# Population-level encoding of task information is stronger in the adolescent compared to adult frontal cortex

**DOI:** 10.1101/2024.10.31.621384

**Authors:** Madeline E. Klinger, Hongli Wang, Lung-Hao Tai, Albert J. Qü, Mei Murphy, Linda Wilbrecht

## Abstract

Adolescence is commonly claimed to be a ‘sensitive period’ for brain development, but it is not clear how experience at this stage can have such strong impacts. We hypothesized that enhanced sensitivity to experience during adolescence may manifest as stronger encoding of task-related information in the dorsomedial prefrontal cortex (dmPFC). To enable optical access to task-related layer 2/3 neural activity in the developing mouse dmPFC, we imaged mice under a 2-photon microscope while they learned an auditory go/no-go task. We found adolescent mice (postnatal day P30-45) learned the task to criterion faster than adult mice (P60-75). When we compared neural activity in expert mice with comparable performance between the two age groups, we found that a similar fraction of single cells encoded task variables in the two groups. However, task information could be better decoded from the adolescent dmPFC population activity than the adult, even when we controlled for differences in head-fixed running. Adolescents also showed greater noise correlation than adults and shuffling to remove this noise correlation suggested noise correlation contributed to gain of function in adolescent compared to adult brain. We suggest a working model for adolescent brain function in which greater noise correlation supports greater capacity for distributed encoding of information driving, for better or for worse, increased sensitivity to experiences at this stage of life.

---

There is strong interest in an adolescent neural plasticity with a particular focus on frontal lobe development, but there is no consensus on why experiences at this age can be so formative. In studies of rodents in the laboratory, a large body of research has shown that adolescent experience in the third to fifth week of life (∼P21-40) can significantly impact adult brain and behavior (1–4). The details of the experiments vary widely, but it is not uncommon to find that experiences in the adolescent period have a greater impact than experiences in adulthood on later cognitive and decision-making (1, 3–5), affective and social behavior (6–10), and even sensory discrimination (11). These animal studies may also explain why adolescence may be a period for increased risk of developing mental illnesses and substance use disorders (4, 12–14).

Many different neural circuits and brain regions could be affected by adolescent experience, but the prefrontal cortex (PFC) is of particular interest due to its protracted development and role in cognitive control and affective regulation (1, 3, 4, 14, 15). Despite growing confidence in enhanced sensitivity to adolescent experience at the behavioral level, we do not yet have community consensus on the mechanism for the cellular-level ‘capture’ of this information. Higher spine formation rates, higher spine density, changes in the balance of long range and local connectivity, and changes in inhibitory and/or neuromodulatory signals have all been identified as possible candidates (15, 16). However, metrics of *ex vivo* long-term potentiation are weaker in the adolescent frontal cortices compared to adult (17). Hope for a consensus mechanism is also complicated by questions about differences in cortical regions, circuits, cell types and layers and the ideal stage in development to investigate for each.

We and others have argued that we can use studies of the primary auditory and visual cortices as a ‘Rosetta stone’ for understanding how the neurons of the frontal cortices might regulate a putative sensitive period during adolescence (15, 16, 18). An extensive body of literature in the primary sensory cortices demonstrates that sensitive period plasticity in Layer 2/3 of the neocortex is regulated by an increase in GABAergic inhibitory neurotransmission on pyramidal (PYR) neurons (19–21). If this increase is blocked, for example by *Gad65*-knockout, then mice fail to enter a naturally occurring period of receptive field plasticity in the binocular cortex (22), but this plasticity can be later rescued by administration of a GABAA receptor agonist, specifically through activation of the α1 subunit (23, 24). More recently, transplant studies suggest GABAergic neurons themselves convey critical factors. Transplant of embryonic inhibitory neurons into the visual cortex of young mice was shown to induce a later sensitive period, once the implanted neurons matured to the age of typical sensitive period onset (25, 26).

An emerging body of work suggests the maturation of GABAergic inhibition also occurs in the dorsomedial frontal cortex (dmPFC) of the mouse during early adolescence which could regulate neural plasticity in this region (15, 18, 27, 28). mIPSC recordings reveal an increase in the strength of inhibitory neurotransmission on to L2/3 and L5 IT-type PYR in *ex vivo* brain slices of the dmPFC from P25 to P40 regulated by puberty onset (27). Optogenetic stimulation of parvalbumin positive interneurons (PV INs) reveals these neurons develop increasing capacity to silence L2/3 PYRs from ∼P23 to P60 (29). Studies of the mEPSC/mIPSC ratio in L2/3 PYRs suggest there is an inflection between P27-28 and P31-33 when the E:I ratio reaches adult levels and tonic inhibition onto L2/3 PYR decreases (27). Moreover, recent chemogenetic and pharmacogenetic studies suggest adolescent, but not adult, inhibition of mPFC PV INs or thalamo-prefrontal projections can disrupt pyramidal neuron firing properties and have detrimental effects on cognition (28, 30). Based on these data and classic results from developing sensory systems, we speculated that the increase in GABAergic inhibition in L2/3 of the dmPFC L2/3 around ∼P30 could transiently enhance function in this frontal region (15, 27).

Other cellular-level changes may also contribute to enhanced plasticity in the frontal cortices during adolescence. For example, spine density and turnover on L2/3 and L5 PYR both decrease throughout medial frontal cortices between P29-P60 (29, 31–36), perhaps mediated by microglia (29, 37). Additionally, dopamine neuron activity (38), dopamine axon innervation (39, 40) and dopamine receptor density mature in the medial frontal cortices during adolescence and may contribute to changes in interneuron function (41). Mesocortical dopamine axons are also more plastic in face of ‘experience’ in adolescents than in adults as evidenced by increased bouton formation following optogenetic stimulation (42).

Changes in dopamine, GABAergic neurotransmission, and spine density are likely happening in conjunction (4, 15, 43). In a simple model when these are all occurring in the dmPFC in adolescence, enhanced plasticity might lead to enhanced functions, such as enhanced learning which can occur in some contexts (2) or enhanced encoding (the subject of this manuscript). However, adolescent brain changes are more often associated with immaturity in function, which has often been judged to be inferior to adult function (44–47). For example, adolescent behavior has been shown be more exploratory, noisy, or error prone (2, 48, 49) and primary auditory cortical regions are known to be noisier than adults (44). Computational explorations of learning that model differences in spine density and spine pruning have recently been used to explain why adolescents may perform worse than adults (50). However, immaturity in behavioral function may also be explained by a learning-performance gap driven by enhanced exploratory behavior in adolescents (4). It is possible that information may be encoded in the adolescent frontal cortices that does not always translate to behavior.

To deepen our understanding of adolescent frontal function at the cellular level, we sought to empirically measure how well neurons in the dmPFC encode information in adolescence compared to adulthood, ideally when behavioral performance was not different between groups. Since adolescent mice are smaller than adults, we chose a head-fixed imaging environment in which mice did not have to carry headgear. We then chose an auditory go/no-go task because it has been shown to be rapidly learned (4) and sensitive to disruption of the dmPFC by muscimol and optogenetic silencing of dmPFC neurons that project to the striatum (51, 52). We chose to compare neural activity in mice learning the task in the P30 and P60s because the documentation of GABAergic inhibitory development specifically in L2/3 of dmPFC shows marked increase and change in the E:I ratio at ∼P30 (27).

These results of these experiments could help arbitrate between three possible hypotheses. If there is an adolescent period where experience has magnified impact on frontal circuits, then we would predict that dmPFC may show greater capacity for encoding task-related information in the P30s than in the P60s. If, on the other hand, the mature adult brain achieves a more optimal state for encoding, we might instead expect that the adult dmPFC would exhibit more robust encoding of task information compared to adolescents. We also considered a null hypothesis that despite differences in connectivity and plasticity, the adolescent and adult dmPFC may encode information about the task comparably despite support by different connectivity configurations.

## Results

### Adolescent mice learned an auditory discrimination task faster than adults

We trained male and female adolescent (P30-44) and young adult (P60-74) mice (CaMKII-tTa^+/-^ / tetO-GCaMP6s^+/+^) that expressed calcium indicator GCaMP6s in cortical pyramidal neurons in a head-fixed go/no-go task. Prior to training, all mice underwent surgery for implantation of a cranial window centered over the left dmPFC, encompassing the medial secondary motor area (M2) and most dorsal aspect of the anterior cingulate (Cg1). We recorded calcium transients in L2/3 PYR during each training session in all animals and obtained high-quality imaging data in 10/14 adolescent (6/9 expert) and 11/24 adult (5/15 expert) mice (**Fig. 2A, B; Supplemental Table 1**). Adolescent and adult expert mice completed a comparable number of trials per session (**Supplemental Fig. 1**).

We chose a simple go/no-go auditory discrimination task because it could be rapidly learned within a 14-day period (**Fig. 1A**). Training began with habituation to a 7 kHz pure tone ‘go’ cue (‘cue 2’), which signaled the availability of a 4 μl water reward. Once mice consistently licked the waterspout in response to the go cue (typically 1-2 sessions), we introduced a 14 kHz ‘no-go’ cue (‘cue 7’), which required inhibition of licking for the 3 s response window; licking to a ‘no-go’ cue triggered a 10 s time-out. Discrimination between the ‘go’ and ‘no-go’ cues was measured by *d’*, the ratio of z-scored hit rate to z-scored false alarm rate. Through pilot studies, we set a criterion for ‘task mastery’ at *d’* = 1.8, corresponding to a false alarm rate of 0.7 given perfect go cue responses (hit rate modeled at 0.99). To facilitate rapid learning, if an animal sustained performance at *d’* > 1.8 for 50 trials on the initial 2 cues, the next pair of ‘go’ and ‘no-go’ cues were immediately introduced. This was repeated until all four pairs were included in a session.

**Figure 1:**
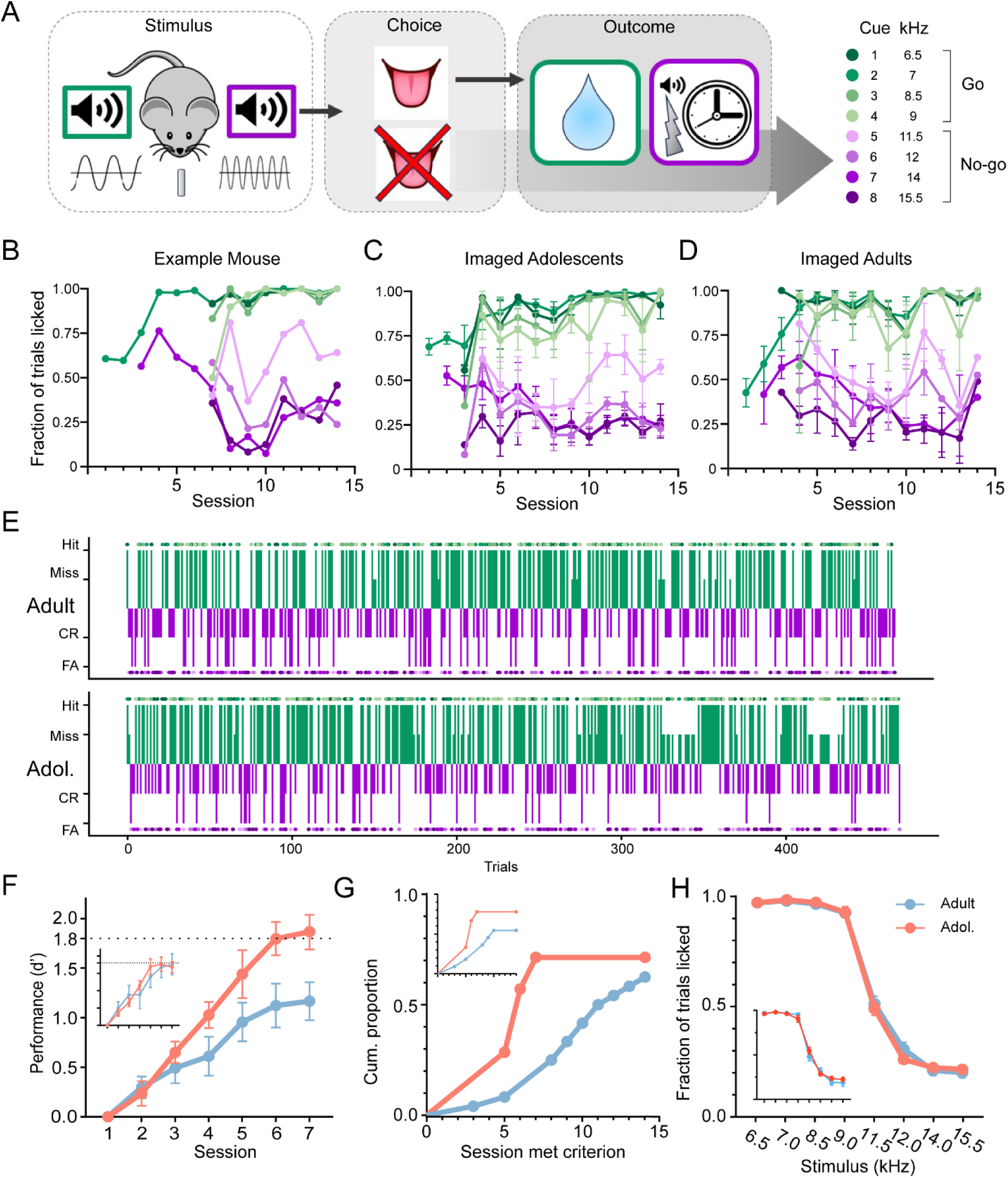
Adolescents learn a go/no-go task more quickly than adults but show comparable performance once expert. **A)** Schematic of task. **B)** Example learning trajectory from one adolescent mouse, displaying lick rates to all cues present in each session. Colors specified in A). This mouse reached criterion at session 6 and expert level performance at session 8. **C)** Learning trajectories from all imaged adolescent mice (n = 6). Error bars represent SEM. **D)** Learning trajectories from all imaged adult mice (n = 5). Error bars represent SEM. **E)** Example sessions of adult (top) and adolescent (bottom). Go and no-go trials are plotted in green and purple, respectively. Individual cues are represented by dots in different shades of green and purple. Four different trial types, hit, false alarm (FA), correct rejection (CR), and miss, are shown on the y-axis. **F)** Performance by session for expert animals (n = 9 adolescent, n = 15 adult); dotted line = criterion; inset = animals with high-quality imaging data who became expert by the end of 14 days. **G)** Cumulative proportion of animals that have reached criterion by session; inset = animals with high-quality imaging data who became expert by the end of 14 days. **H)** Performance on the task was comparable when focusing on expert mice from both groups. Showing data from experts in all late phase sessions in main panel; inset = late phase performance sessions from experts where imaging quality was suitable for subsequent single cell encoding and population level decoding analyses). See also Supplemental Table 1.

**Figure 2:**
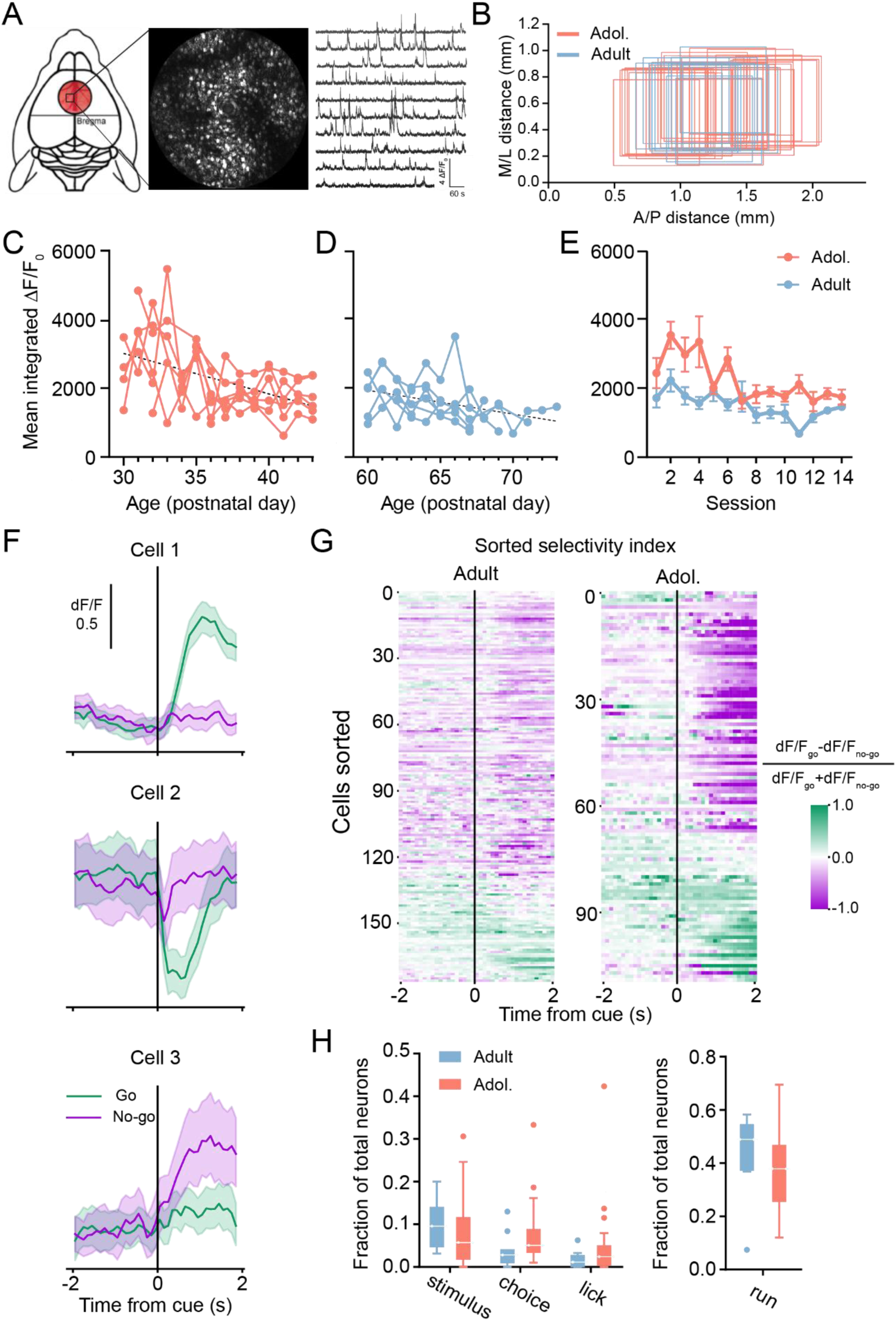
Similar proportions of neurons encode task variables in adult and adolescent dmPFC. **A)** Schematic of imaging area, example field of view (FOV), and example Ca2+ traces. **B)** Imaging FOV locations for sessions in (F). **C)** Mean integrated fluorescence from all cells in all adolescent animals across sessions; individual lines = individual animals; dotted line = linear regression. **D)** Mean integrated fluorescence from all cells from all adult animals across sessions; individual lines = individual animals; dotted line = linear regression. **E)** Mean of (C) and (D) with SEM in error bars. **F)** Mean fluorescence traces from three example cells, aligned to the sound cue and averaged across go (green) and no-go (purple) cues. The shaded area shows the 95% confidence interval. **G)** Heat map of cue selectivity for the cells of an example session from adult (left) and an example from adolescent (right) as a function of time. Cue selectivity was calculated as the normalized difference between mean fluorescence traces from go and no-go trials. Green and purple shadings indicate preference for go and no-go cues, respectively. Cells were sorted by the center-of mass of their selectivity traces. **H)** Fraction of total neurons imaged that encoded task variables: stimulus (go/no-go), choice (lick/no lick), and overall licking throughout the trial. Box shows median, first and third quartile; whiskers represent quartiles ± 1.5IQR. See also Supplemental Table 4.

The ‘complete task’ therefore consisted of 8 pure tone cues that tiled the mouse auditory spectrum (‘go’ cues: 6.5, 7, 8.5, 9 kHz; ‘no-go’ cues: 11.5, 12, 14, 15.5 kHz; cues are numbered 1-8 from 6.5-15.5 kHz for convenience in Figs. 1-4). Animals learned to lick in response to all four go cues quickly, while gradually suppressed licking when no-go cues were presented (**Fig. 1B-D**). Mice were considered ‘expert’ when they achieved an average *d’* > 1.8 on all 8 cues for at least one session. Mice trained for one hour each day for 14 consecutive days, or until mice had performed above criterion on all 8 cues for 3 consecutive days. Expert adult and adolescent animals typically performed around 500 trials per session with comparable performance (**Fig. 1E**).

Examining performance of ‘expert’ mice in the first 7 sessions, before performance plateaued, we found that adolescent mice learned at a faster rate than adults (mixed-effects ANOVA, interaction of age × session: F(6,131) = 2.672, *p* = 0.018; **Fig. 1F**). As a metric of learning efficiency, we quantified the number of training sessions required for mice to reach our *d’* > 1.8 criterion for all 8 cues. Of the animals trained, 10/14 adolescent and 15/24 adult mice reached criterion on at least one session within 14 days (one adolescent reached criterion on the first cue set but was unable to learn the complete task). Notably, adolescents reached criterion sooner than adults (adolescent median = day 6, adult median = day 11.5; *p* = 0.017, Gehan-Breslow-Wilcoxon test; **Fig. 1G**). Note that for calcium imaging analysis, we only examined data from adolescent and adult mice that reached expert status for all cues (see data shown in inset in all panels).

At the end of the 14-day training period, a similar fraction of mice from both age groups had reached the *d’* > 1.8 criterion on all 8 cues (9/14 adolescent and 15/24 adult mice; *χ*2 = 0.012, df = 1, *p* = 0.912). We next compared late performance (after session 5) in the subset of ‘expert’ mice from both groups, defined as mice maintaining *d’* > 1.8 on all 8 cues for at least one session. To this end, we compared licking rates to each of the 4 ‘go’ and 4 ‘no-go’ cues, combining sessions across 9 expert adolescents and 15 expert adults (n = 42 expert adolescent sessions, n = 33 expert adult sessions). We found that expert mice in both groups showed comparable licking to ‘go’ cues (adult mean lick rate = 0.96 ± 0.01, adolescent mean lick rate = 0.97 ± 0.01) and comparable inhibition of licking to ‘no-go’ cues (adult mean lick rate = 0.31 ± 0.03, adolescent mean lick rate = 0.30 ± 0.02), as determined by two-way ANOVA (main effect of age: F(1,73) = 0.034, *p* = 0.855; main effect of cue: F(7, 511) = 945.3, *p* < 0.001; age × cue interaction effect: F(7,511) = 0.757, *p* = 0.624, **Fig. 1H**). Thus, although adolescents learned the task faster than adults, when experts of both groups were studied, their licking performance was comparable for the 4 ‘go’ and 4 ‘no-go’ cues.

### Similar proportions of adult and adolescent dmPFC neurons encode task information across learning at single-cell level

We next analyzed neural activity using calcium imaging in L2/3 in the expert adolescent and adult groups (**Supplemental Table 2**). We decided to focus our analysis only on mice that became experts by the end of 14 days to isolate neural differences more relevant to age differences in encoding while removing variation related to performance differences. Imaging data were sampled from comparable regions of the dmPFC in both groups (**Fig. 2A, B**).

First, we plotted overall neural activity levels from expert mice over the two weeks. We calculated the mean area under the curve from the ΔF/F0 traces of all cells imaged for each session as a proxy measure of neural activity for that day. Adolescent activity by this metric was higher than adult activity in the early P30s, then decreased to adult-like levels by mid P30s. Linear regressions revealed a negative slope indicating a decrease in activity in both adolescent (slope = -117.8, *r^2^* = 0.245, *p* < 0.001; **Fig. 2C**) and adult groups (slope = -68.53, *r*^2^ = 0.153, *p* = 0.005; **Fig. 2D**). Mixed effects ANOVA showed a main effect of age group (F(1,11) = 16.18, *p* = 0.002; **Fig. 2E**) and session (F(3.4,24.1) = 3.26, *p* = 0.034).

We next sought to investigate the extent to which dmPFC encoded task-related variables and whether expert adolescent dmPFC represented task variables differently from expert adult dmPFC when behavioral performance was comparable. At the single-neuron level, some recorded cells responded differently in go or no-go trials (**Fig. 2F**). To quantify the different responses, we calculated the selectivity index as the normalized difference between the two mean traces. During the period following the cue onset, neurons from both adults and adolescents exhibit go and no-go selectivity (**Fig. 2G**). To further examine the single-neuron encoding of task-relevant variables, we constructed a mixed linear regression model with coefficients representing task variables: stimulus (‘go’ vs. ‘no-go’), and animal choice (lick vs. no lick). Because animal motion is encoded throughout the cortex (52, 53), we also included terms to capture neural representation of animal licking and running. We applied the regression model to imaged expert sessions (adult n = 13 sessions, 92.8 ± 12.4 cells per session; adolescent n = 28 sessions, 127.7 ± 9.4 cells per session) and evaluated the proportion of neurons with significant coefficients for each variable. We found no main effect of age group, but a small interaction between task variable and age group (mixed effects ANOVA, effect of task variable: F(3, 114) = 141.2, *p* < 0.001; effect of age group: F(1, 38) = 0.185, *p* = 0.670; interaction effect: F(3, 114) = 2.965, *p* = 0.035; **Fig. 2H**). In adults vs. adolescents reported respectively, we found 10.2% ± 1.6% vs. 7.9% ± 1.4% (mean ± SEM) of imaged neurons encoded stimulus and 3.7% ± 1.0% vs. 7.1% ± 1.2% encoded choice. Only a very small percentage (1.6% ± 0.5% in adults vs. 4.7% ± 1.5% in adolescents) encoded licking, and more than a third of all recorded neurons encoded running (44.2% ± 4.1% in adults vs. 36.9% ± 2.6% in adolescents). Together, these data suggest that the fraction of single cells strongly encoding task information is comparable in performance-matched adolescents and adults.

### Task variables can be decoded with greater integrity from adolescent brain compared to the adult brain

A growing body of research suggests that information represented by the frontal cortices may not be linearly encoded in single neurons but instead is distributed in neural activity at the population level, with individual cells exhibiting mixed selectivity for internal and external variables (54–62). To determine if the representations of the task-related information differed in adolescent and adult dmPFC population activity, we trained a support vector machine to predict the identity of the ‘go’ or ‘no-go’ cues based on population calcium activity during early learning (sessions 1-5, before most animals had achieved expert status) and late learning (sessions 6+ in which animals achieved expert status) sessions.

In the early learning phase using data only from the group of animals that would become experts, we found that the stimulus identity could already be decoded at a level higher than chance (see **Methods**)(adult 0-1 s, *p* = 0.027; adolescent 0-1 s, *p* < 0.001; adult 1-2 s, *p* = 0.026; adolescent 1-2 s, *p* < 0.001, Wilcoxon rank sum test, FDR_BH corrected, cell number capped at 80 cells per session per mouse, **Supplemental Table 4**). This indicated that both adults and adolescents rapidly gained a representation of the task without extensive training. During early learning sessions, stimulus identity decoding accuracy showed no difference between age groups (two-way ANOVA, effect of age group, F(1,932) = 1.36, *p* = 0.243; effect of time (within trial), F(1,932) = 82.1, *p* < 0.001, **Fig. 3A**). The average decoding accuracy in the time window 1-2 s after cue onset improved as the number of neurons included in the decoding model increased (two-way ANOVA, effect of size, F(1,232) = 28.1, *p* < 0.001, **Fig. 3B**; total cells adult: 129.0 ± 14.7 cells per session; adolescent: 153.4 ± 11.5 cells per session), consistent with previous reports (53). Meanwhile, the modulation effect of cell population size did not differ between age groups (two-way ANOVA, effect of age, F(1,232) = 0.0143, *p* = 0.905, **Fig. 3B**).

**Figure 3.**
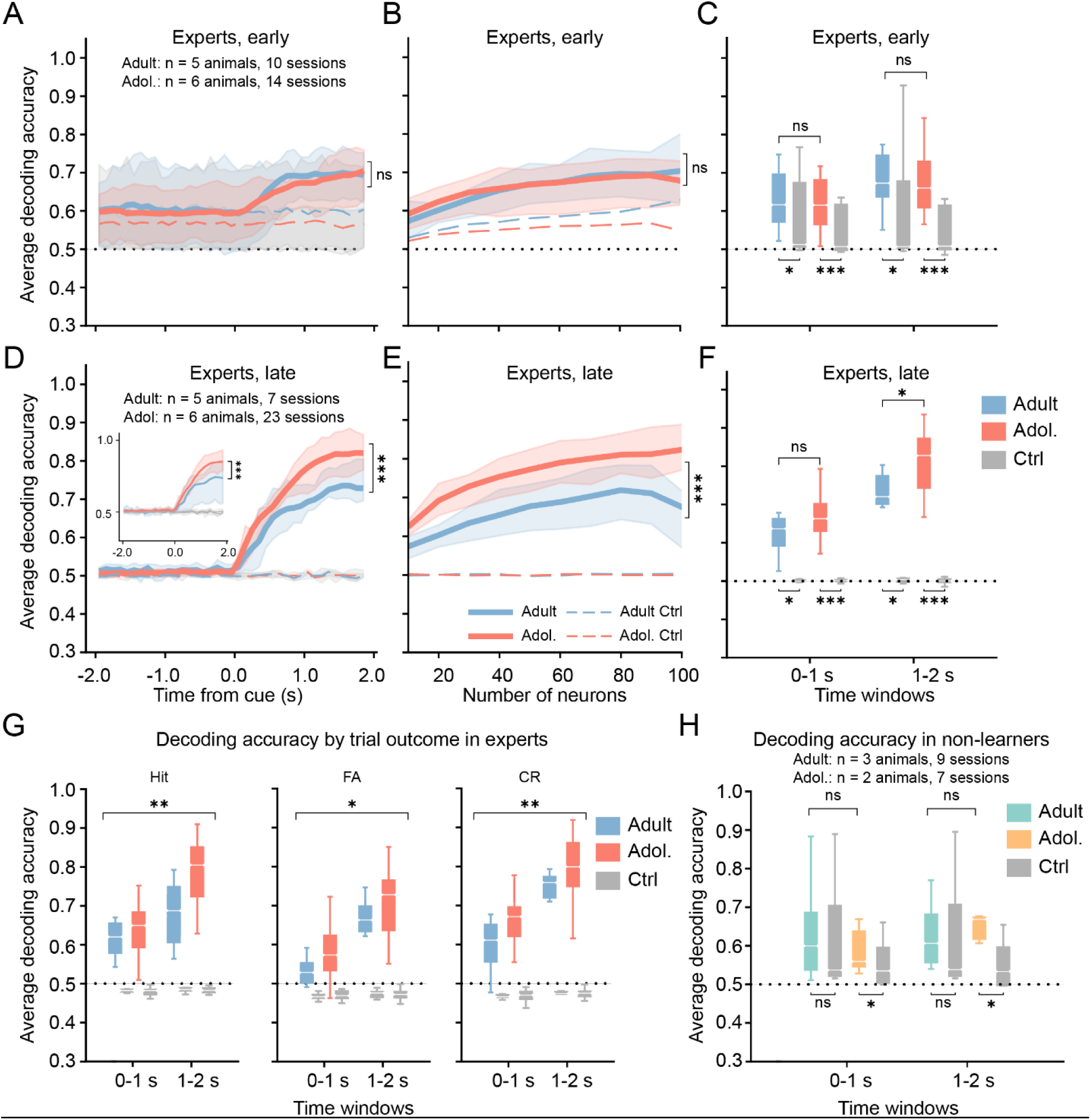
Representation of task stimuli in dmPFC PYR neurons was higher in adolescent experts than in adult experts, but only after reaching expert status. **A)** Decoding accuracy of stimulus as a function of time aligned to cue onset for early learning stage (sessions 1-5). Data are from the subset of mice who would later be categorized as experts (Experts, early). (See (H) for data from those who did not become experts). Sample was limited to 80 cells per session to match sampling from the two age groups. Red: adolescent, blue: adult, gray: shuffled dataset. Note that the decoding accuracy in the shuffled data exceeded 50% due to the greater proportion of ‘go’ relative to ‘no-go’ trials during the early learning sessions (see **Methods**), a factor that was not present at the late learning stage. **B)** Average decoding accuracy between 1-2 s after cue onset as a function of the number of neurons used in the decoding model for expert animals in the early learning stage. **C)** Average decoding accuracy of data shown in (A) binned over 0-1 s and 1-2 s after cue onset in sample capped at 80 neurons per session. Box shows median, first and third quartile; whiskers represent quartiles ± 1.5 IQR. **D)** Decoding accuracy of stimulus as a function of time aligned to cue onset for late learning phase (sessions 6-14, **Supplemental Table 3**). After mice were expert (Experts, late), an effect of age group emerged: more information could be decoded from the adolescent data than the adult data. This result held even when we limited our sample to the first three sessions after criterion was reached in each individual (a control for days of experience at criterion) (Supplemental Table 3). **E)** Average decoding accuracy between 1-2 s after cue onset changed as a function of the number of neurons used in the decoding model again in the late learning stage. **F)** Average decoding accuracy of data shown in (D) binned over 0-1 s and 1-2 s after cue onset in sample capped at 80 neurons per session. More information could be decoded from the adolescent than the adult data in the 1-2 second window. **G)** Average decoding accuracy of expert adults (blue) and adolescents (red) at the late learning stage, separated by trial outcomes Hit, false alarm (FA) and correct reject (CR). A significant effect of age group could be seen for all outcome types. **H)** Same analysis as (C) but for non-learners. ns: not significant; *: *p*<0.05; **: *p*<0.01; ***: *p*<0.001. See also **Supplemental Figure 2** and **Supplemental Tables 4-6**.

**Figure 4.**
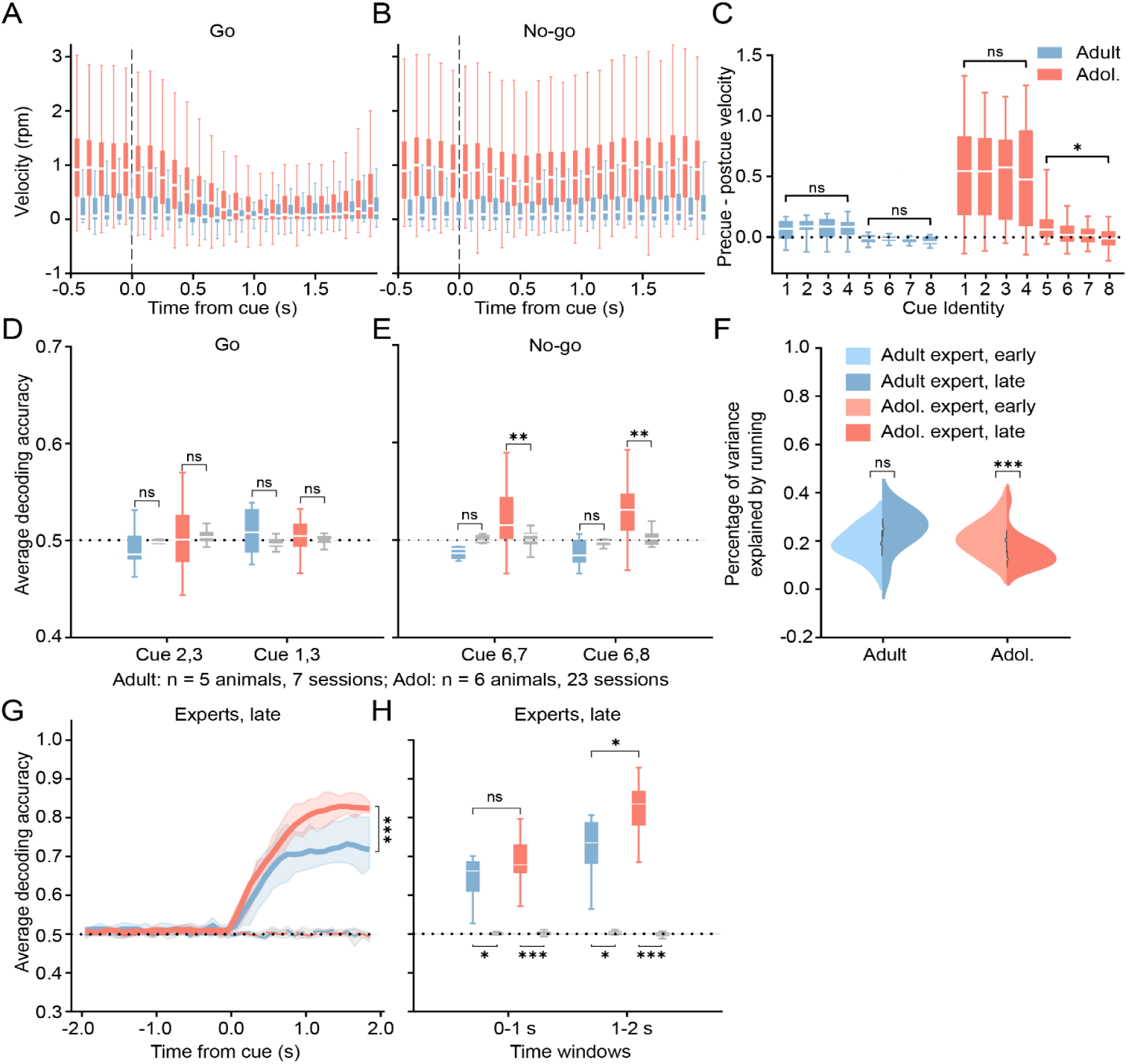
Better decoding in adolescents cannot be explained by differences in running behavior. **A)** Distribution of running velocities for expert animals segregated by stimulus type for ‘go’ trials. Box and whisker plot shows median, first and third quartile, and ±1.5 IQR **B)** Same analysis as (A) for ‘no-go’ trials. **C)** Difference in running velocity (precue mean - postcue mean) by cue. Box shows median, first and third quartile; whiskers show min to max. **D)** Average decoding accuracy in 1-2 s after cue onset for cues 2 vs. 3 (left) and cues 1 vs. 3 (right) with ensemble size of 80 at the late learning stage. Blue: adult experts at the late learning stage; Red: adolescent experts at the late learning stage. **E)** Same analysis as (D) for cues 6 vs. 7 (left) and cues 6 vs. 8 (right). **F)** Distribution of the percentage of variance explained by running speed in the calcium activity for adult (left) and adolescent (right) at the early (light blue and light red for adult and adolescent, respectively) and late (dark blue and dark red for adult and adolescent, respectively) learning stages. **G)** Decoding accuracy of stimulus as a function of time aligned to cue onset for late learning stage. In this analysis, the neural activity used in the decoding model was adjusted to remove the possible running effect. Analysis was capped for both age groups at 80 cells per session. Red: adolescent, blue: adult, gray: shuffled dataset. **H)** Average decoding accuracy of data shown in (G) binned over 0-1 s and 1-2 s after cue onset. Analysis was capped for both age groups at 80 cells per session. ns: not significant; *: *p*<0.05; **: *p*<0.01; ***: *p*<0.001. See also Supplemental Tables 5-7.

We also plotted the averaged decoding accuracy in two time bins: 0-1 s and 1-2 s after cue onset. These data showed again that the stimulus information can be decoded from the calcium activity in both age groups, yet there were no significant differences between age groups during this early learning stage (0-1 s, *p* = 0.977, 1-2 s, *p* = 0.661, Mann-Whitney test, **Fig. 3C**).

We next examined stimulus identity decoding accuracy during late learning sessions, after these same animals had become experts. At this later phase of training, decoding accuracy after cue onset was again significantly higher than chance in both age groups and adolescent decoding accuracy was now significantly higher than adult (two-way ANOVA, effect of age group, F(1,116) = 36.6, *p* < 0.001; effect of time, F(1,116) = 2.68×10^3^, *p* < 0.001. Cells per session capped at 80 for both groups; **Fig. 3D**), indicating that the stimulus variables were more accurately represented in the neural activity of the adolescent mice. To rule out the possibility that more adolescent sessions from later stages of training may bias the decoding accuracy, we ran the same analysis on a subset of data including only the first three sessions after the performance met the criteria. The decoding accuracy of adolescent group still exceeded that of adult **(Fig. 3D**, inset, *p* < 0.001), suggesting that the effect was not due to longer times at criterion. Again, the decoding accuracy depended on the number of neurons included in the decoding model (two-way ANOVA, effect of age group, F(1,282) = 72.6, *p* < 0.001; effect of size, F(1,282) = 130, *p* < 0.001, **Fig. 3E**; adult: 92.8 ± 12.4 cells per session; adolescent 127.7 ± 9.4 cells per session). Finally, we again plotted the averaged decoding accuracy in two time bins: 0-1 s and 1-2 s after cue onset. In both of these time bins, decoding accuracy was higher than chance (adult 0-1 s, *p* = 0.016; adolescent 0-1 s, *p* < 0.001; adult 1-2 s, *p* = 0.016; adolescent 1-2 s, *p* < 0.001, Wilcoxon rank sum test, FDR_BH corrected, **Fig. 3F**), and the average decoding accuracy in the adolescent group remained significantly higher than that of adults in the 1-2 s time bin (0-1 s, *p* = 0.069; 1-2 s, *p* = 0.037, Mann-Whitney test, **Fig. 3F**).

In the next analysis, we used the same decoding model to decode the animals’ choice in the current trial (lick or no lick), as well as the type of outcomes of the trial (hit, false alarm, or correct rejection) in the 0-1 s and 1-2 s post cue periods. We found we could accurately decode choice in the current trial and the three different kinds of outcomes with higher than chance accuracy in both the early and late learning phases. Again, in the late learning phase but not early learning phase, we found decoding accuracy was significantly stronger for adolescents’ data when compared to adult (**Supplemental Fig. 2**). Taken together, these results suggested an enhanced representation of task-related variables in adolescent population activity compared to adults. The fact that this age difference took training time to develop and reflected different outcomes suggests that it was mediated by learning and/or task experience.

Last, to determine if the stimulus representation is modulated by reward, we plotted the decoding accuracy separated by different trial outcomes (hit, false alarm, and correct rejection). The results showed an overall significant age group effect across trial types (two-way ANOVA, effect of age, F(1,84) = 13.3, *p* < 0.001; effect of trial, F(2,84) = 6.94, *p* = 0.001; **Fig. 3G, Supplemental Table 5**), suggesting an enhancement of information representation in the dmPFC of adolescent mice that was consistent across all possible outcomes.

Importantly, the difference in the decoding accuracy between age groups was absent in non-learners, mice of both age groups that had extended training but failed to learn the task. In these non-learners, in early and late sessions combined, the decoding accuracy was not significantly higher than chance in adult, and only marginally higher than chance in adolescents (adult 0-1 s, *p* = 0.652; adolescent 0-1 s, *p* = 0.031; adult 1-2 s, *p* = 0.568; adolescent 1-2 s, *p* = 0.031, Wilcoxon rank sum test, FDR_BH corrected, **Fig. 3H, Supplemental Table 6**).

### Enhanced decoding in adolescents cannot be explained by differences in running in the task

Given that over one third of the imaged neurons encoded animal running, differences in running could be a major influence on decoder results. We examined running speeds in the [-2s, 2s] window around cue onset during both ‘go’ and ‘no-go’ trials from adolescent and adult expert sessions and found that running speed was significantly modulated by stimulus type and age, with adolescents running faster than adults and both ages pausing after cue onset on ‘go’ trials (two-way ANOVA, effect of stimulus type: F(1,38) = 34.45, *p* < 0.001; effect of age: F(1,38) = 15.23, *p* < 0.001, interaction: F(1,38) = 15.92, *p* < 0.001; **Fig. 4A, B**). We next evaluated running speed modulation after different cues within each stimulus category specifically from the expert sessions used for imaging analyses. We observed minimal running modulation in response to ‘no-go’ cues in adolescents, and a slight increase in speed in adults only (adolescent: *p* = 0.069, adult: *p* < 0.001, one-sample t-test against 0 modulation; **Fig. 4B**).

While both adults and adolescents significantly modulated their running in response to ‘go’ cues (adult: *p* = 0.024, adolescent: *p* < 0.001, one-sample t-test against 0 modulation; **Fig. 4A**), average adolescent modulation was 7.33-fold greater in magnitude (**Fig. 4C**). Therefore, we considered the possibility that the observed decoding difference might emerge from running-related neural activity, rather than representation of the stimulus.

To examine and control for the role of running-related neural activity, we took three approaches. First, we attempted to control running by comparing decoding of cues for which running and running modulation was comparable. We built a new decoding model to discriminate between pairs of within-category stimulus cues (i.e., cue 2 vs. cue 3). Low or ‘at-chance’ decoding results for these within-category comparisons might be expected because the ability to discriminate between individual cues under the same ‘go’ or ‘no-go’ category was not beneficial in the behavioral task. We found that, while the decoding model was not able to discriminate between any pairs of ‘go’ cues in either age group (example pair in **Fig. 4D**; see **Supplemental Table 7** for full data), the decoding model was able to classify ‘no-go’ cue identity at a level significantly higher than chance for multiple pairs of ‘no-go’ cues using adolescent data (**Fig. 4E**).

Meanwhile, the decoder was only ‘at chance’ for all pairs of ‘no-go’ cues when trained with adult data (example cue pair in **Fig. 4E**; see **Supplemental Table 7** for full data). This result suggests that adolescent dmPFC had a more specific representation of individual ‘no-go’ cues than the adult brain and this difference was not likely to be driven by encoding of movement, since the running speed and modulation was comparable for many of the decodable ‘no-go’ cue pairs (**Fig. 4C, E**).

As a second method to explore the role of running on task representation, we built two multiple linear regression models to fit the recorded calcium activities from the task-related variables. One model used the choice, stimulus, and running speed as predictors. The other used choice and stimulus but not running (see **Methods**). By comparing two models, we can estimate how dependent the calcium activity is on running speed (and running modulation) by calculating the amount of variance explained by the latter. The results showed an overall low contribution of running speed in the early and late learning stages for both age groups (**Supplemental Table 8**). Moreover, we found that running explains less variance in adolescent group in later learning, after mice become expert (adult: n = 11 sessions for early learning, n = 12 for late learning; *p* = 0.103; adolescent: n = 16 for early learning, n = 28 for late learning; *p* < 0.001; Mann-Whitney U test, **Fig. 4F**), suggesting the enhanced decoding accuracy in adolescents that emerges with learning was not paralleled by an increase in running-related neural activity.

Lastly, based on the multiple linear regression results, we calculated adjusted ΔF/F0 to subtract the contribution of running from the calcium dynamics. The same decoder was then fit to the adjusted calcium data. Since it is possible that the neurons in the dmPFC encoding task-related information project to downstream brain areas to modulate running, this treatment could potentially affect signals that encode task-related information while correlating with downstream running. Nevertheless, the stimulus information could still be decoded with higher accuracy in adolescent compared to adult (two-way ANOVA, effect of age group: F(1,116) = 44.58, *p* < 0.001; effect of time: F(1,116) = 2.91×10^3^, *p* < 0.001; **Fig. 2G**. Mann-Whitney U test: 0-1 s, *p* = 0.158; 1-2 s, *p* = 0.007; **Supplemental Table 9, Fig. 2H**).

Taken together, these three attempts to control for the effects of running suggest that the higher decoding accuracy in the adolescent dmPFC is likely more than just an artifact of running or running modulation. Additionally, the comparison on pairs of ‘no-go’ cues further suggests that the representation of task information in the population of L2/3 neurons in the adolescent dmPFC is not only stronger but also more specific than in the adult dmPFC.

### Noise correlation may account for adolescent benefit in information decoding

Previous studies have linked information encoding with the noise correlation in the neural network, with some studies showing noise correlation improves decoding accuracy while others find it decreases decoding accuracy (63–65). Developmental research shows that the adolescent brain is noisier than the adult brain (44), and some of this noise may be due to higher connectivity and higher noise correlation. Therefore, we sought to explore the idea that the enhanced information representation in the adolescent brain is due to enhanced noise correlation. To test this hypothesis, we characterized the pairwise and population noise correlation for expert animals (see **Methods**). The average pairwise noise correlation, estimated by taking the average of the Pearson correlation for every neuron pair, showed a significant effect of learning phase (early vs. late), but not of age group (three-way ANOVA , effect of age group, F(1,125) = 1.96, *p* = 0.164; effect of trial, F(1,125) = 0.729, *p* = 0.395; effect of learning phase, F(1,125) = 6.16, *p* = 0.0144, **Fig. 5A, Supplemental Table 10**). Meanwhile, the population correlation, estimated by taking the first principal component of the PCA performed on the calcium activity of every neuron recorded in a session, showed both significant learning phase and age group effect (three-way ANOVA, effect of age group, F(1,125) = 10.5, *p* = 0.002; effect of trial, F(1,125) = 0.005, *p* = 0.945; effect of learning phase, F(1,125) = 5.87, *p* = 0.017, **Fig. 5B, Supplemental Table 10**).

**Figure 5.**
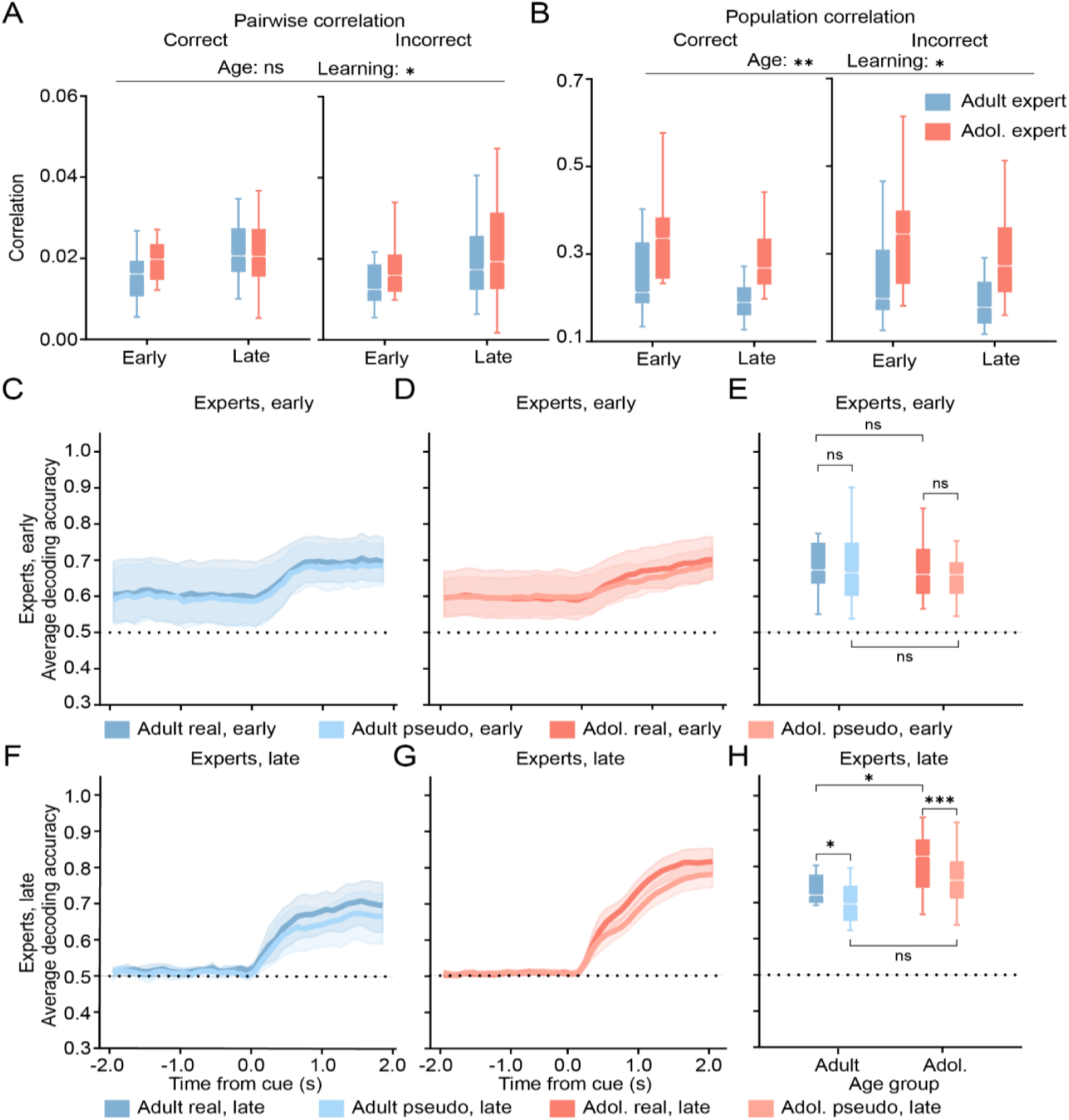
Noise correlation is beneficial to the information encoding in adolescents and adults, and without it, adolescent decoding no longer outperforms adult. **A)** Average pairwise noise correlation in 1-2 s after cue onset for correct trials (left) and incorrect trials (right) differed by learning stage (early vs. late) but not age group. **B)** Population noise correlation differed significantly by learning stage and age group. **C)** Decoding accuracy as a function of time from cue onset for real ensemble (dark blue) and pseudo ensemble (light blue, shuffled to remove noise correlation) for adult experts at the early learning stage. **D)** Decoding accuracy as a function of time from cue onset for adolescent experts at the early learning stage: real ensemble (dark red) and pseudo ensemble (light red). **E)** Average decoding accuracy between 1-2 s after cue onset at the early learning stage for adults (left) and adolescents (right). At this early stage, removing noise correlation did not impact decoding in either age group. **F)** Same analysis as (C) for adults at late learning stage. **G)** Same analysis as (D) for adolescent experts at the late learning stage. **H)** Same analysis as (E) for adult and adolescent experts at the late learning stage. At this late learning stage, removing noise correlation decreased information encoding in both age groups, and eliminated the adolescent decoding advantage over adults. ns: not significant; *: *p*<0.05; **: *p*<0.01; ***: *p*<0.001. See also Supplemental Tables 10-11.

To further examine the noise correlation during the learning process, we constructed a pseudo-ensemble by randomly shuffling the trial identity within trial types (hit, false alarm, and correct rejection) for every neuron independently to maintain the signal correlation while removing the noise correlation across neurons (**see Methods**). For adult and adolescent experts at the early learning stage, we found that removing the noise correlation in the neural ensembles did not affect the decoding accuracy (adult real, *p* = 0.193; adolescent real, *p* = 0.058, Wilcoxon rank sum test, **Fig. 5C-E, Supplemental Table 11**) and did not create or reveal any age group differences (real, *p* = 0.661; pseudo, *p* = 0.747, Mann-Whitney U test; **Supplemental Table 11**). However, removing the noise correlation at the late/expert stage resulted in a significant reduction in decoding accuracy for ‘go’ and ‘no-go’ stimuli in both adolescents and adults (adult: *p* = 0.016; adolescent: *p* < 0.001, Wilcoxon rank sum test; **Fig. 5F-H, Supplemental Table 11**), which is consistent with previous reports for this region (66). Notably, the decoding accuracy in the pseudo-ensemble of both age groups was not different (*p* = 0.077, Mann-Whitney U test, **Fig. 5H, Supplemental Table 11**), indicating that noise correlation contributed to the adolescent advantage over the adult age group in encoding information. Taken together, these data suggest that noise correlation in the dmPFC emerges with experience, noise correlation contributes to decoding success, and higher noise correlation in adolescents compared to adults may contribute to greater decoding accuracy in this age group.

## Discussion

Our study intended to explore functional encoding of information in the adolescent versus the adult brain. We reasoned that the encoding of simple task variables could be a metric of capacity for plasticity and function that we could use as a dependent variable to compare the adolescent and adult dmPFC that would overcome issues of learning-performance gap. To enable access to the functional activity of single neurons in the developing mouse, we used a rapidly learned head-fixed go/no-go task in male and female mice starting at P30 and P60 and training for up to 14 days.

We anticipated that results from these experiments could help arbitrate between competing ideas about adolescent brain function. We reasoned that if there is enhanced plasticity in L2/3 of the neocortex just after increases in L2/3 GABAergic neurotransmission, then we might see stronger task encoding in the adolescent brain just after P30. However, if the adolescent brain was simply in an ‘immature’ state, then we might see that task encoding was less efficient in P30-45 adolescents than in adults. Finally, we also considered the possibility that despite differences in neural network connectivity and plasticity, adolescents and adults might encode task information comparably. We refer to this as the null hypothesis.

Our results reveal some support for the enhanced adolescent plasticity hypothesis and some support for the null hypothesis. We found that adolescent mice learned a head-fixed go/no-go task more quickly than adult mice, but with time, expert mice in both groups reached a comparable level of performance which could facilitate head-to-head neural encoding comparison. Examination of neural data from our performance-matched ‘expert’ mice in both age groups revealed that the fraction of individual neurons that encode task variables in L2/3 dmPFC was comparable across ages. At the population level, however, decoders trained with adolescent data in later training stages showed greater decoding accuracy compared to adults. Adolescent trained decoders also could discriminate between individual ‘no-go’ cues, while decoders trained with adult data could not. These data suggest that information about the task stimuli was richer and more specific in the adolescent than adult brain, even with an adjusted dataset that removed the running modulation from the calcium activity. Finally, noise correlation was higher in the expert adolescent mice compared to the expert adult mice later in training.

When we removed this noise correlation, the adolescent decoding advantage was no longer apparent, suggesting noise correlation contributed to the adolescent information encoding advantage seen in later training phases. The fact that adolescents learned the task faster than adults is consistent with the concept of enhanced capacity for plasticity in adolescence. However, it may also be surprising because adolescent prowess in ‘frontal’ tasks is unexpected. The literature is dominated by studies that find that adult rodents and nonhuman primates can outperform adolescents in laboratory tasks that tax adolescent executive functions including behavioral inhibition (67, 68), delay discounting (69, 70), instrumental extinction and contingency degradation (71, 72), working memory (17, 41, 44, 73), and cognitive flexibility (41, 74). However, it is notable that adolescent rodents have been observed to learn faster or to reverse faster than adults on other frontal-dependent tasks (2, 75–77) and can be seen to perform similarly to adults in some metrics (71, 77). We can perhaps reconcile these divergent findings by considering that adolescent prowess may be limited to specific forms of learning and/or to specific developmental windows (2). It also may have been critical that our mice were habituated to a running wheel and operant water before training and that we trained our mice alone. Adolescent mice trained with ‘peers’ may (like human teens) show a peer effect on their sensitivity to reinforcement and/or negative outcomes (78). We also introduced new cue pairs when mice were still learning and did not wait for asymptotic performance on previously introduced cues. This might have enhanced sensitivity to cues and reduced reliance on stimulus-insensitive one-back strategies that can increase with age and task experience in mice (79). We may also consider that adolescents at P30 are also lighter in weight and their motivation to work for food or water may be heightened when they are in a phase of rapid body growth. They may also be more resilient to the effects of recent surgery. These factors could explain why adolescents were more efficient at learning the go/no-go task in our study, but also may explain adolescent prowess or resilience under challenging circumstances ‘in the wild.’ Therefore, these may be ‘features’ of adolescent biology rather than ‘a bug’ in our experimental design, where treatment of the mice was as fully-matched as possible.

Importantly, we do not depend on early behavioral performance for our main conclusions. Our major findings come from expert adolescent and adult mice in later training when both groups met learning criteria for all cues and exhibited comparable behavior (**Fig. 1D**). This allowed us to study neural activity without confounding differences in performance. Here we focused on the L2/3 of dmPFC, a region that receives input from the mediodorsal thalamus (one way to define prefrontal areas). Based on previous muscimol, optogenetic, and chemogenetic manipulations we were confident that the dmPFC region was critical to successful performance in a go/no-go licking task (51–53) that could be rapidly learned by our adolescent and adult mice.

Previous studies comparing cellular-level neural activity in the awake, behaving adolescent and adult prefrontal cortex brain are rare. Of the existing studies there are some results that are consistent, some orthogonal, and none, we would argue, conflicting. Studies of the dorsolateral prefrontal cortex (dlPFC) in rhesus macaques performing working memory tasks found that adolescent and adult responses to cues were comparable, except on trials where the monkey aborted the trial, which was more common in adolescents (48). When monkeys completed the trials, differences did emerge in neural activity during the delay period, when the adolescent dlPFC maintained a less active response than the adult to the now-absent cue (48). The adolescent monkey dlPFC neurons also showed stronger responses to ‘distractor stimuli’ than the adult (48, 80). Recent work in rodents has also appeared in preprint recording via prism in a deeper medial PFC region of the medial prefrontal cortex in a CS+/ CS-discrimination task (81). In this study, activity in adolescents was found to be lower for CS-cues but comparable or larger for CS+ cues, and adolescent mice achieved more hit trials and licked more on CS+ trials compared to adults. Although this pattern was not observed in our data, it may parallel our observation of a performance advantage in adolescents during our head-fixed go/no-go task.

Given these behavioral differences in performance from past studies, it was therefore particularly valuable to be able to examine neural encoding in the brain at adolescent and adult mice when performance was comparable. Our multiple linear regression analysis revealed that a similar fraction of neurons encoded task-related variables in adolescents and adults (**Fig. 2H**). These data suggest that despite anatomical and physiological differences between the adolescent and adult brain, they can represent information in single cells similarly. The responses of these individual dmPFC cells were highly complex, but these fractions were comparable to previous reports that 5-10% of adult L2/3 dmPFC neurons are stimulus encoding and 2-5% encode choice (52, 53, 59, 62). Our data were also in concordance with existing reports of cortical activity from widespread areas encoding motor actions (52, 53, 82); we found that more than a third of neurons in our mice were identified by multiple linear regression as encoding running, but less than 5% encoded licking. There are also PFC studies that find the fraction of single cells encoding task variables that are much higher, ranging from 16 to 25% in adults for stimulus, choice, or outcome (52).

We did not see age differences between adolescent and adult mice in single cell encoding estimates. However, in another study, representation of overall task variables in a CS+/CS-licking task was greater in mPFC in adolescents compared to adult mice (65.8% vs 47.7% respectively), and activity specifically in response to CS-was lower in adolescents than adults (81). This study was different from ours in that it examined all cortical layers and was centered on a more ventral region of the medial PFC using a prism. As mentioned above, behavioral performance in the two age groups was also different (81).

Our final and main variable of interest was decoding accuracy. This metric is used to assess the representation of task variables distributed across the entire population of neurons in the frontal cortices (56, 60, 83–88). To compare how much task information was encoded by the population of neurons recorded in the dmPFC of adolescent and adult mice, we implemented a support vector machine decoder, which was trained on the entirety of session imaging data thus able to discern differences between trials that emerge from population-level activity that are not discriminable through our multiple linear regression model. We found that our decoder was able to discriminate well between stimulus types, action choice, and trial outcome at above chance levels (**Fig. 3D-F**, **Supplement Fig. 2**). Moreover, at several time points in the [0,2] second window after cue onset, task information was more readily decoded from adolescent dmPFC data than from adults.

We next examined data taken in earlier stages of learning from the same expert mice. We found that we could decode task information from these data but there were not yet differences between age groups. These results suggest that some aspect of this adolescent gain of function is due to learning or other experiences.

While these data support the hypothesis that the adolescent brain may show enhanced plasticity that leads to more robust information encoding, we also considered the possibility that this result could simply be due to differences in motor behavior between the two groups (which also may reflect stronger learning) (**Fig. 4A-C**). To avoid this confound, we trained our decoder to discriminate between cues within the same ‘go’ or ‘no-go’ category for which running was comparable (**Fig. 4D, E**; **Supplemental Table 6**). Though decoder performance in this application is less successful than discrimination between ‘go’ and ‘no-go’ cues, we could still decode ‘no-go’ cue identity statistically above chance from adolescent data at multiple time points in the [0,2] second window after cue onset. Adult data did not support decoding above chance, suggesting that the adolescent neural population represented ‘no-go’ cue information with higher fidelity or specificity than the adult dmPFC, independent from running differences.

Interestingly, we did not observe the same specificity of encoding for ‘go’ cues. Previous work in mice has shown that optogenetic or muscimol inactivation of dmPFC enhances false alarm rate in a go/no-go task (51, 52), and human fMRI studies of go/no-go tasks have demonstrated greater frontal cortical activation during ‘no-go’ trials (89–91). We thus speculate that decoding between ‘no-go’ cues may be attributed to greater dmPFC engagement when mice are required to inhibit an action.

Our results show that noise correlation was higher in the neural population of adolescent dmPFC, consistent with higher connectivity due to higher spine density (50, 92). To determine if noise correlation contributes to the populational representation of task-related information, we shuffled the trial labels of recorded calcium activity within the same trial types independently for every neuron. With this method, noise correlation of the constructed pseudo-ensemble was removed, yet the signal representation remained intact (63, 65). As expected, the representation of the stimulus was largely maintained in the pseudo-ensemble, as evidenced by the high decoding accuracy of the stimulus in both age groups (**Fig. 5C-H**). Comparison of decoder performance in the original and pseudo condition revealed that noise correlation contributed to the success of decoder trained on both adult and adolescent data. Notably, removing the noise correlation resulted in a comparable level of decoding accuracy across age groups (**Fig. 5H**), suggesting that the noise correlation supports the enhanced information encoding in adolescents. While potentially unexpected, the direction of this result has been previously documented (63, 65, 93). Thus, this analysis supports the idea that higher connectivity and/or correlated activity between neurons may facilitate encoding in the neocortex (65, 67, 93) and may explain adolescent prowess in our study.

In sum, our data support the enhanced plasticity in adolescence hypothesis and aspects of the null hypothesis. We found that at the level of individual cells, adolescents and adults encoded task information comparably, but at the population level, the capacity to encode information was greater in adolescents. One might speculate that the greater capacity for encoding task information may have enhanced the efficiency of learning in the adolescent mice. However, the encoding advantage in adolescents was observed only after they reached expert-level behavior, not during early learning. Further experiments may be needed to sample the period of rapid learning in adolescents in dmPFC and other brain regions to evaluate how this superior encoding emerges. It could also be important to test how adverse events or drug-related cues are encoded to understand how adolescence may be a sensitive period for effects of adverse experience and development of substance use disorders.

Manipulations of age and inhibitory and dopaminergic mechanisms, as well as spine structural plasticity, may also be used in future experiments to test the mechanisms underlying noise correlation and changes in population encoding capacity in the frontal cortices. Future work on underlying biological mechanisms and their time course will be a valuable future direction to help us understand why adolescent experience is so formative. Understanding the intertwined role played by changes in GABA, spine pruning and stabilization, gonadal hormones, and the maturation of dopamine and other modulators (43, 47, 50, 94–100), may even elucidate ways to sculpt adult cortical plasticity to facilitate educational, psychiatric, or neurological interventions. It should be noted, however, that we cannot assume enhanced capacity for encoding in adolescence is solely beneficial. Enhanced encoding of task events in the dmPFC of the adolescent brain may reflect compensation for some other aspects of adolescent function that are weaker (101), and this difference in encoding may not uniformly benefit all aspects of neural or behavioral function.

In conclusion, we have presented evidence that adolescents learn an auditory go/no-go task faster than adults, and that task information is more accurately decoded from neural population activity in L2/3 dmPFC of the adolescent brain compared to that of the adult, arguably due to enhanced noise correlation. While further work is required to more clearly delineate the cellular and synaptic mechanisms regulating these functional differences, we posit that enhanced encoding of experience over the time scale of days in adolescence could underlie the lasting impacts of adolescent experience into adulthood.

## Resource Availability

Data collected and analyzed in this study and code generated for analyses will be made available upon publication in an open science repository.

## Author Contributions

Madeline E. Klinger: Investigation and data curation (lead); methodology, formal analysis, software, and writing - original draft (equal). Hongli Wang: Investigation and data curation (supporting); methodology, formal analysis, software, and writing - original draft (equal). Lung-Hao Tai: Methodology and software (equal). Albert Qü: Software (equal). Mei Murphy: Investigation (supporting). Linda Wilbrecht: Conceptualization, funding acquisition, supervision, and writing - reviewing and editing (lead).

## Acknowledgements

We thank Adi Mizrahi, Benne Praegel, Bruno Averbeck, Michael Stryker, Na Ji, Hillel Adesnik, Nuria Vendrell-Llopis, Pierre-Marie Dominique Garderes, and Wan Chen Lin for discussion, advice, and comments. We thank Katrina Wong, Katrina Manaloto, Samantha Jackson, and Bailey Donnelly for assistance with experiments and manuscript preparation. This work was supported by a grant from the UCSF UC Berkeley Schwab Dyslexia and Cognitive Diversity Center Innovation Fund (to L.W. and M.K.).

## STAR Methods

### Key Resources Table

| REAGENT or RESOURCE | SOURCE | IDENTIFIER |
| --- | --- | --- |
| Deposited data |  |  |
| Raw and analyzed data | This manuscript | *Repository TBD*<br>*Data currently available on request* |
| Experimental models: Organisms/strains |  |  |
| Mouse: CaMKII-tTA: B6.Cg-Tg(Camk2a-tTA)1Mmay/DboJ | The Jackson Laboratory | RRID: IMSR_JAX:007004 |
| Mouse: TRE-GCaMP6s line G6s2: B6;DBA-Tg(tetO-GCaMP6s)2Niell/J | The Jackson Laboratory | RRID:IMSR_JAX:024742 |
| Software and algorithms |  |  |
| Suite2P | Pachitariu et al. [99] | https://suite2p.readthedocs.io/en/latest/ |
| NoRMCorre | Pnevmatikakis [100] | https://github.com/flatironinstitute/NoRMCorre |
| Preprocessing pipeline | This manuscript | https://github.com/wilbrecht |
| Behavioral and imaging analysis pipeline | This manuscript | https://github.com/wilbrecht |

### Experimental Model and Study Participant Details

We bred transgenic mice in-house, crossing tetO-GCaMP6s mice (https://www.jax.org/strain/024742) and CaMKIIa-tTA mice (https://www.jax.org/strain/007004). We used male and female mice heterozygous for the CaMKIIa-tTA transgene and homozygous for the tetO-GCaMP6s transgene. All mice were weaned on postnatal day (P)21 and housed in groups of 3-4 same-sex siblings on a 12:12 h reversed light:dark cycle (lights on at 22:00 h). Experimental mice belonged to two age cohorts: adolescent (P30-44) and adult (P60-74); experimental mice were housed alongside nonexperimental littermates. All animals were given access to food and water *ad libitum* prior to experiment start, after which they were maintained on *ad libitum* food and 1 mL water per day until experiment end. All procedures were approved by the Animal Care and Use Committee of the University of California, Berkeley and conformed to principles outlined by the NIH Guide for the Care and Use of Laboratory Animals.

## Method Details

### Surgeries, Handling, and Head-fixed Behavior

Male and female mice (P28 or P58) were deeply anesthetized with 5% isoflurane (vol/vol) in oxygen and placed in a stereotaxic frame with ear bars (Kopf Instruments) on a heating pad for the duration of surgery. Anesthesia was maintained at 1%–2% isoflurane during surgery. An incision was made along the midline of the scalp and a 3 mm diameter circular craniotomy performed over the anterior cingulate area of the left hemisphere medial prefrontal cortex (coordinates relative to Bregma: A/P: -0.5 to 2.5 mm, M/L: -2.0 to 1.0 mm). Stacked glass coverslips of 3 mm and 5 mm diameter (Electron Microscopy Sciences) bonded with UV optical adhesive (Norland) were implanted over the craniotomy and sealed with dental cement (C&B Metabond® Quick Adhesive Cement, Parkell, Inc.). A stainless steel off-centered circular head bar (custom design, eMachineShop) was affixed above the cranial window. Mice were given subcutaneous injections of meloxicam (10 mg/kg) during surgery and 24 and 48 h after surgery.

Four nights before surgery, access to water was restricted. The following morning, mice were placed in an arena with food and *ad libitum* water available from mounted lick spouts to habituate mice to water-seeking behavior in a novel environment. After 4-6 hours, mice were removed from the arena and handled for at least 5 minutes before undergoing habituation to a running wheel placed in their home cage for 15 minutes. Mice underwent pre-training, handling, and running wheel habituation for a total of 3 days, receiving at least 1 mL water each day.

Water was returned to the home cage after the third day of pre-training, and surgery was performed the following day. Due to rapid bone regrowth over the cranial window in adolescent animals, imaging began as soon as possible after surgery, following approval from the Animal Care and Use Committee. Thus, 24 hours after surgery, mice were administered analgesic, handled for 5 minutes, then habituated to the experimental arena for 15 minutes. Mice were placed on an acrylic running wheel rotating about the Y-axis and head-mounted to a custom steel bracket mounted above the wheel. Mice received access to water during this time through a lick spout. After habituation, mice were returned to the home cage. Water was removed from the home cage that evening. The next morning the experiment began. Weight was monitored daily.

In the go/no-go task, mice were trained to discriminate between 4 ‘go’ and 4 ‘no-go’ auditory cues (500 ms each) by licking to ‘go’ cues and inhibiting licking to ‘no-go’ cues. Water restricted mice began training on this task 48 hours after surgery, at P30 or P60. Mice trained one session per day for at most one hour. Mice were head-fixed atop a disc running wheel with access to a lick spout, which was connected to a capacitance sensor. Water rewards (2 μL) were delivered by gravity drip regulated through a solenoid valve controlled by TTL pulses. On ‘direct delivery’ trials mice received one reward, and on ‘hit’ trials, two water rewards separated by 100 ms were administered for a total of 4 μL. Auditory stimuli were delivered through speakers (Bohlender & Graebener, Neo3-PDR Planar Tweeter) placed on either side of the wheel. The task was administered through custom scripts written in MATLAB.

On the first day of training, all mice began the experiment with at least 50 ‘direct delivery’ trials. In these trials, a 500ms ‘go’ cue (7 kHz pure tone, ‘cue 2’) indicated the availability of a water reward, and mice had 3 s to respond by licking. Mice were immediately given a double reward (4 μL water) upon licking; if they did not lick, a single reward (2 μL water) was given at the end of the lick interval. Water delivery initiated an inter-trial interval (1-2 s, drawn from an exponential distribution with tau = 2), after which mice must refrain from licking for an additional 1 second before subsequent cue onset. Direct delivery trials continued until mice regularly began to lick in response to the auditory cue, receiving a double reward. After 20 consecutive double-rewarded trials, direct delivery ended; from that point on, mice had 3 s after cue onset to lick, and water (double reward) was delivered only if mice licked (‘hit’ trial). If mice did not lick after 3 s, no reward was given, and the trial was labeled a ‘miss’. After a further 20 hit trials, a 500 ms ‘no-go’ cue (14 kHz pure tone, ‘cue 7’) was introduced at a 1:9 ratio of no-go:go cues. In response to a ‘no-go’ cue, mice must refrain from licking for 3 s in order to complete a “correct reject” trial and progress to the inter-trial interval. If mice licked (‘false alarm’ trial), incorrect action was signaled by a 500ms white noise burst and punished with a 10s timeout before progressing to the inter-trial interval. We noted during pilot sessions that introducing the first ‘no-go’ cue at a 1:1 ratio with the ‘go’ cue dissuaded most animals from participating in the task at all; thus, we employed a progressive strategy to introduce the first ‘no-go’ cue first at a 10% frequency, increasing the frequency as tolerated by each animal to 20%, 30%, and finally 50%. By presenting the ‘no-go’ cue with increasing frequency, animals gained positive reinforcement from the more-frequent ‘go’ cue, and thus maintained motivation to continue engaging with the task. If mice behaviorally disengaged from the task upon increasing the occurrence of ‘no-go’ cues, the proportion of ‘no-go’ cues was reduced with the goal to encourage consistent licking to ‘go’ cues.

Once mice achieved tolerance of 1:1 no-go:go cues, we began tracking their performance using the metric *d’*. A rolling *d’* value was calculated over the past 50 trials as *d’* = z(hit rate) - z(false alarm rate). We used the MATLAB function *norminv*, which calculates the inverse of the standard normal cumulative distribution function, to approximate a z-score. Because the *norminv(x)* function approaches infinity and negative infinity as *x* approaches 1 and 0, respectively, the upper bound of the hit rate and false alarm rate was set to 0.99 and the lower bound was set to 0.01. We determined a threshold of *d’* = 1.8, corresponding to a 100% hit rate and 70% false alarm rate (i.e., 30% correct ‘no-go’ trials and 100% correct ‘go’ trials). If mice sustained performance of *d’* > 1.8 for 20 consecutive trials, we introduced an additional cue set (‘go’: 6.5 kHz, ‘cue 1’; ‘no-go’: 15 kHz, ‘cue 8’) alongside the first ‘go’ and ‘no-go’ cues for a total of 4 different tones at a 1:1:1:1 ratio. Similarly, after sustaining *d’* > 1.8 for 20 consecutive trials after at least 50 trials of these 4 cues, the next cue set was introduced (go: 8.5 kHz, ‘cue 3’; no-go: 12 kHz, ‘cue 6’) in equal proportion to existing cues; after sustaining *d’* > 1.8 for 20 consecutive trials after at least 50 trials of these 6 cues, the final cue set was introduced (go: 9 kHz, ‘cue 4’; no-go: 11.5 kHz, ‘cue 5’) in equal proportion to existing cues. Finally, after sustaining *d’* > 1.8 for 20 consecutive trials after at least 50 trials of all 8 cues, 8 probe cues were introduced at a 1% frequency each. These cues approximated ‘go’ and ‘no-go’ cues in that they preceded a 3 s response window prior to the 1-2 s inter-trial interval, but they did not yield reward or trigger a timeout, regardless of the mouse’s response. These probe trials were omitted from current analysis due to insufficient trials. In order to be considered to have learned the complete task and to be at ‘expert’ level, mice must have achieved an average *d’* > 1.8 over at least one session with all cues and probes present. We defined the period of ‘early learning’ to include sessions 1-5 and ‘late learning’ to include sessions 6 and after. Mice were trained for 14 consecutive days; experiments were ended earlier if mice either completed 3 consecutive sessions with all cues and probes or completely disengaged from the task for 3 consecutive sessions. Mice achieved various levels of completion, and learners who became expert completed comparable numbers of trials per session (see **Supplemental Figure 1, Supplemental Table 3**).

### Two-photon imaging

We performed calcium imaging of layer 2/3 of the dorsomedial prefrontal cortex (dmPFC), defined here as the area comprising most dorsal aspect of the anterior cingulate area and the medial 1 mm of the supplementary motor cortex (M2) (centered coordinates relative to Bregma: A/P: +0.5 to 2.0 mm, M/L: 0 to 1 mm, D/V: -0.2 to -0.35 mm; **Fig. 2B**). Mice were imaged through a cranial window using a commercial two-photon microscope (Bruker) controlled by PrairieView software, while they were head-fixed and performing the task Excitation light was provided by a Ti:Sapphire laser (Mai Tai, Newport Spectra-Physics) tuned to 910 nm; the beam was focused on layer 2/3 through a 20x water-immersion objective (NA 1.0, Olympus). Images were acquired through spiral scanning at 8 Hz, with a resolution of 256 x 256 pixels and frame diameter of 650 μm (0.39 pixels/μm). Windows were visually screened prior to training each day and were only imaged during that session if cells were clearly visible (see summary in **Supplemental Table 1**). Registration of imaging data was completed either in Suite2P (102) or NoRMCorre (103). Images were aligned with one round of non-rigid registration (in 64 x 64 pixel chunks) and up to five rounds of rigid registration. Images were manually evaluated for sufficient motion correction and quality prior to cell detection and were discarded if motion was still present or cells were insufficiently visible. All cell detection was completed in Suite2P, where each ROI was manually validated to be classified as a cell. Sessions with fewer than 50 cells were omitted from imaging analyses. Validated sessions were neuropil subtracted, and baseline fluorescence (F0) was determined for each cell as follows: we calculated the mean and standard deviation for fluorescence values in a rolling 200-frame window, then set a threshold = mean + 1.5 standard deviations and calculated the mean of all frames in that window under the threshold. ΔF/F0 was calculated for each cell for each frame and used as the basis for all imaging analyses.

### Quantification and Statistical Analysis

To obtain trial-averaged activity traces associated with cue onset, we first aligned ΔF/F(t) traces based on their timing relative to each instance of the cue and then took the mean across traces. To estimate confidence intervals, we performed a bootstrapping procedure, as follows. For N instances of a particular event, N traces were resampled randomly with replacement and then averaged over 1000 iterations, in order to approximate the sampling distribution for the mean.

Upper and lower bounds of the confidence interval were then estimated as percentiles of this distribution. Cue selectivity was calculated as the difference between cue-aligned trial-averaged traces from trials in which the go versus no-go cue was presented, divided by the sum of the two traces.

Statistical analysis was performed either through custom MATLAB scripts (MathWorks Inc.), custom Python scripts using the *SciPy* package, or in GraphPad Prism (GraphPad Software Inc.). Our decoder was implemented through the Scikit-learn package for Python, using *sklearn.svm.SVC*. Data are reported as mean +/- standard error of the mean, unless otherwise noted.

To determine how fluorescent signals may relate to choices and outcomes, we used multiple linear regression adopted from previous literature (66, 104):

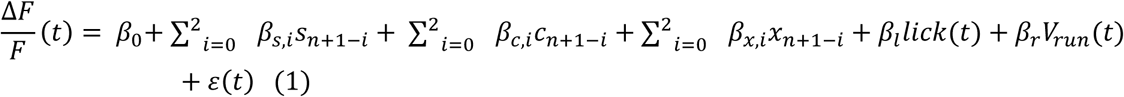

where 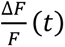 is the fractional changes in fluorescence at time t in trial n; *s* represents stimulus identity (‘go’ vs. ‘no-go’), *c* represents the animal’s action choice (lick vs. no lick), and *x* represents interactions between *s*, and *c*; subscripts *n+1*, *n*, and *n-1* represent the next trial, the current trial, and the previous trial, respectively; *β*_0_*,…, βl* are the regression coefficients; *lick*(*t*) takes the value of 1 if the animal licked at time t and is otherwise 0; *Vrun* (*t*) is the running speed of the animal at time *t*; and *ɛ*(*t*) is the error term. For each session, the regression coefficients were determined by fitting the equations to data using python package *statsmodels*. Equations were fit in 100 ms time bins spanning from -2 to 2 s relative to cue onset, using mean 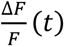 within the time bins. For a given predictor and an ROI, if the regression coefficients were significant (*p* < 0.01) for at least 3 consecutive or 10 total time points, the ROI was considered significantly modulated by the predictor. To summarize the results, for each predictor, we calculated the proportion of ROIs in which the regression coefficient was significantly different from zero (*p* < 0.01). To determine if the proportion was significantly different from chance, we performed a chi-square test against the null hypothesis that there was a 1% probability that a given predictor was mischaracterized as significant by chance in a single session. To determine if the proportion of different age groups were significantly different from each other, we performed a chi-square two sample test against the null hypothesis that the two groups were drawn from the same binomial distribution.

To determine how running speed modulates calcium activity, we built two multiple linear regression models:

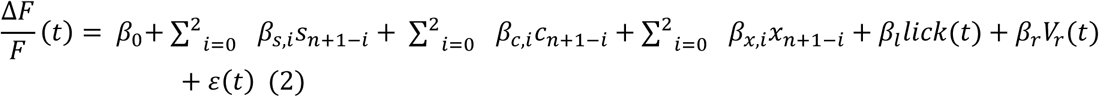

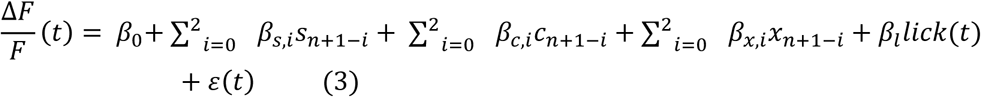

The percentage variance of calcium activity explained by running speed is obtained by taking the difference of the variance explained by the first model and the second model:

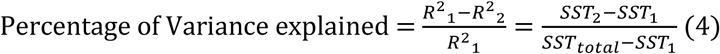

To decode trial information from our calcium imaging neuronal data, we implemented a support vector machine using the Scikit-learn package for Python, *sklearn.svm.SVC*. Each trial was classified by stimulus type (‘go’ or ‘no-go’), animal choice (lick or no lick), or trial outcome (hit, false alarm, or correct reject). 50% of trials from each session were used as a training dataset, and the remaining 50% of trials comprised the test dataset (a 70-30 split of trials for the training and test datasets was also tested; this did not change the conclusion. Data not shown). This support vector machine projects our data into an n-dimensional space, where n = number of neurons in the recorded session, and aims to optimize a hyperplane between data points of different categories. The hyperplane was optimized through maximization of the margin between points of different categories. Decoder performance on the test dataset was compared to a control dataset, in which category labels are shuffled across trials for every neuron independently to approximate chance level. This process was performed 20 times using unique training vs. test datasets to cross-validate the decoding result. Fluorescence data was interpolated every 50 ms, and decoder performance was analyzed on every point of this interpolated data within the [-2s, 2s] window centered at cue onset. To determine the decoding accuracy for each trial type separately, after training the model on the same training and test dataset as previously described, trials in the test dataset were grouped based on the trial outcomes (hit, false alarm, and correct rejection) and the prediction accuracy on each trial outcome was calculated within each group. For decoding accuracy with a fixed number of neurons, the neurons included in the model are randomly chosen without replacement from the whole population to build the decoding model. The final decoding results are obtained by taking the average of 10 random picks to overcome the sampling variability.

Two-way ANOVA was performed to determine the effect of age and time on the decoding accuracy time series. For head-to-head comparisons, average decoding accuracy was taken between two one-second intervals, 0-1s and 1-2s after cue onset. Mann-Whitney tests were used for comparisons between learning stage and age groups. Wilcoxon rank sum tests were used for comparisons between actual data and shuffled controls. Correction of multiple comparisons was performed across similar comparisons with false discovery rate, Benjamini– Hochberg procedure (FDR_BH).

To eliminate the possible running effect on the recorded calcium activity, we calculated adjusted 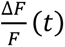 as the following equation based on prior multiple linear regression. Then we trained the decoding model on the adjusted 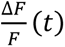.

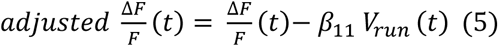

To characterize the effect of noise correlation on decoding accuracy, we constructed a pseudo-ensemble to remove the noise correlation in the population calcium activity. The trial labels were shuffled across trials within the same trial outcome (hit, false alarm, and correct reject) to maintain the signal correlation. The shuffling was performed independently for every neuron.

For each neuron pair, we calculated pairwise noise correlation by taking the Pearson correlation between the df/f at each time points within correct or incorrect trials separately, as

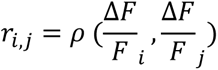

where *i*, *j* are the index of the neurons. The results were averaged across neuron pairs, and 1-2s after cue onset to obtain the average pairwise correlation.

For the population activity of correct or incorrect trials, we quantified the population noise correlation by performing principal component analysis (PCA) on each time point and taking the percentage of variance explained by the first principal component (63). The population correlation was then averaged across trial type and 1-2s after cue onset to obtain the average population noise correlation.

