## Supplemental Information for "Population-level encoding of task information is stronger in the adolescent compared to adult frontal cortex"

**Supplemental Table 1.** Mice trained in this study (in sequential order of training). Highlighted rows represent animals used in imaging analysis.

| Sex | Age Group | # sessions trained | # sessions with high quality imaging data | Behavioral performance achieved | Included or Reason excluded from imaging analysis |
|---|---|---|---|---|---|
| M | Adolescent | 14 | 0 | Learned complete task/Expert | No imaging data |
| M | Adolescent | 14 | 0 | Learned complete task/Expert | No imaging data |
| F | Adolescent | 14 | 0 | Reached cue set #1 | Not expert |
| F | Adolescent | 14 | 0 | Reached cue set #3 | Not expert |
| F | Adult | 11 | 0 | Reached cue set #3 | Not expert |
| M | Adult | 14 | 0 | Learned complete task/Expert | No imaging data |
| M | Adult | 14 | 0 | Reached cue set #4 | Not expert |
| M | Adult | 11 | 0 | Learned complete task/Expert | No imaging data |
| M | Adult | 14 | 0 | Reached cue set #1 | Not expert |
| M | Adolescent | 14 | 13 | Learned complete task/Expert | Included as expert |
| M | Adolescent | 14 | 14 | Learned complete task/Expert | Included as expert |
| M | Adolescent | 14 | 14 | Learned complete task/Expert | Included as expert |
| M | Adult | 14 | 13 | Learned complete task/Expert | Included as expert |
| M | Adult | 10 | 9 | Learned complete task/Expert | Included as expert |
| M | Adult | 9 | 8 | Learned complete task/Expert | Included as expert |
| F | Adult | 13 | 0 | Learned complete task/Expert | No imaging data |
| F | Adult | 13 | 13 | Reached cue set #2 | Not expert |
| M | Adult | 13 | 0 | Learned complete task/Expert | No imaging data |
| F | Adolescent | 13 | 12 | Learned complete task/Expert | Included as expert |
| F | Adolescent | 14 | 5 | Reached cue set #1 | Not expert |
| M | Adolescent | 11 | 7 | Reached cue set #1 | Not expert |
| M | Adolescent | 10 | 10 | Reached cue set #2 | Not expert |
| F | Adolescent | 10 | 8 | Learned complete task/Expert | Included as expert |
| M | Adult | 13 | 3 | Reached cue set #1 | Not expert |
| M | Adult | 12 | 0 | Learned complete task/Expert | No imaging data |
| F | Adult | 13 | 3 | Reached 30% no-go | Not expert |
| F | Adult | 14 | 13 | Reached 30% no-go | Not expert |
| F | Adolescent | 14 | 3 | Learned complete task/Expert | No expert imaging data |
| M | Adolescent | 13 | 12 | Learned complete task/Expert | Included as expert |
| F | Adult | 11 | 0 | Reached 30% no-go | Not expert |
| F | Adult | 14 | 0 | Learned complete task/Expert | No imaging data |
| F | Adult | 14 | 0 | Learned complete task/Expert | No imaging data |
| M | Adult | 14 | 0 | Learned complete task/Expert | No imaging data |
| M | Adult | 14 | 0 | Learned complete task/Expert | No imaging data |
| F | Adult | 9 | 6 | Learned complete task/Expert | No expert imaging data |
| F | Adult | 10 | 10 | Learned complete task/Expert | Included as expert |
| M | Adult | 13 | 6 | Learned complete task/Expert | Included as expert |
| M | Adult | 14 | 3 | Reached cue set #1 | Not expert |

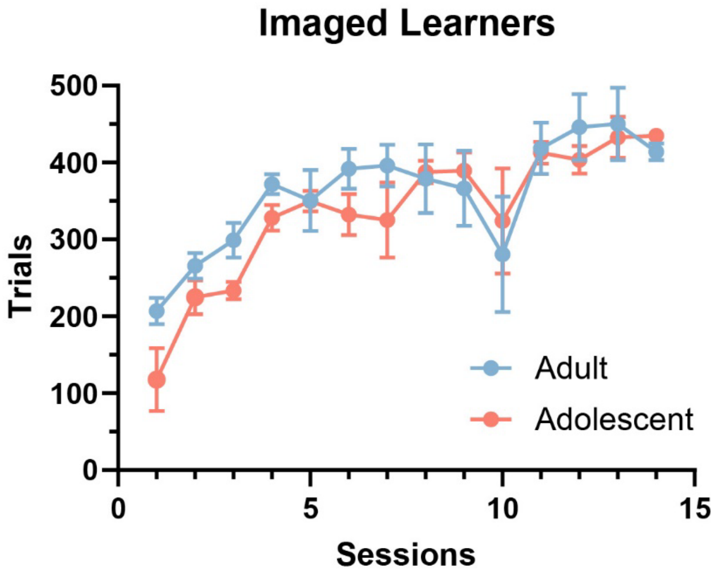


**Supplemental figure 1. Number of trials performed by imaged "expert" animals per session.** n = 6 adolescent, n = 5 adult.

**Session relative to session achieved criterion**

| Animal: | -9 | -8 | -7 | -6 | -5 | -4 | -3 | -2 | -1 | 1 | 2 | 3 | 4 | 5 | 6 |
|---|---|---|---|---|---|---|---|---|---|---|---|---|---|---|---|
| **ADT008** | 0.38 | x | 0.80 | 0.95 | 1.10 | 1.74 | 1.68 | 2.01 | 1.38 | 2.45 | 2.62 | 1.28 | 2.17 | x | x |
| **ADT009** | | | n/a | n/a | 0.45 | 1.17 | 1.56 | 1.54 | 1.08 | 1.76 | 0.82 | 0.78 | x | x | x |
| **ADT010** | | | | | n/a | 0.49 | 1.97 | 2.23 | 2.37 | 2.79 | 2.99 | 3.22 | 1.15 | x | x |
| **ADT028** | | | | | n/a | x* | 0.55 | 1.57 | 1.85 | 2.35 | 2.20 | 3.31 | 3.01 | 2.40 | x |
| **ADT029** | 0.26 | x | x | 1.05 | 1.04 | 1.62 | x | 1.91 | x | 3.03 | x | x | | | |
| **JUV014** | n/a | 0.71 | 0.08 | 0.64 | 0.81 | 1.06 | 1.87 | x | 1.94 | 2.45 | 2.36 | 2.62 | 3.08 | 2.99 | |
| **JUV015** | | n/a | 0.32 | .074 | 0.89 | 2.10 | 1.56 | 1.82 | 1.30 | 2.48 | 2.99 | 2.83 | 2.52 | x | 1.99 |
| **JUV016** | | | n/a | n/a | 0.52 | 1.35 | 1.72 | 2.22 | 1.31 | 2.21 | 2.24 | 3.19 | 2.24 | 2.63 | 2.01 |
| **JUV017** | | | | | n/a | 0.49 | 0.74 | 1.67 | x | 2.32 | 2.38 | 2.75 | 2.75 | x | 1.87 |
| **JUV022** | n/a | n/a | 0.67 | x | 1.87 | 1.06 | x | 1.77 | 1.69 | 1.83 | x | x | x | x | |
| **JUV025** | 0.42 | 1.14 | 1.26 | 1.08 | 2.02 | 1.09 | 1.35 | 1.51 | 1.73 | 2.37 | 2.44 | 2.76 | x | | |

Sessions included in imaging dataset

- Adult, late
- Adolescent, late
- Adult, early
- Adolescent, early

**Supplemental Table 2. Sessions used in imaging analysis aligned to criterion.** Light blue: adult "early" learning sessions. Dark blue: adult "expert" learning sessions. Light pink: adolescent "early" learning sessions. Dark pink: adolescent "expert" learning sessions. Numbers represent overall d' from each session. X indicates imaging or behavioral data was unusable for that session.

**Session relative to session achieved criterion**

| | -12 | -11 | -10 | -9 | -8 | -7 | -6 | -5 | -4 | -3 | -2 | -1 | 1 | 2 | 3 |
|---|---|---|---|---|---|---|---|---|---|---|---|---|---|---|---|
| **ADT008** | 0.38 | x | 0.80 | 0.95 | 1.10 | 1.74 | 1.68 | 2.01 | 1.38 | 2.45 | 2.62 | 1.28 | 2.17 | x | x |
| **ADT009** | | | | | | n/a | n/a | 0.45 | 1.17 | 1.56 | 1.54 | 1.08 | 1.76 | 0.82 | 0.78 |
| **ADT010** | | | | | | | | n/a | 0.49 | 1.97 | 2.23 | 2.37 | 2.79 | 2.99 | 3.22 |
| **ADT028** | | | | | | | | n/a | x* | 0.55 | 1.57 | 1.85 | 2.35 | 2.20 | 3.31 |
| **ADT029** | | n/a | x | 0.26 | x | x | 1.05 | 1.04 | 1.62 | x | 1.91 | x | 3.03 | | |
| **JUV014** | | | | | 0.71 | 0.08 | 0.64 | 0.81 | 1.06 | 1.87 | x | 1.94 | 2.45 | 2.36 | 2.62 |
| **JUV015** | | | | | n/a | 0.32 | .074 | 0.89 | 2.10 | 1.56 | 1.82 | 1.30 | 2.48 | 2.99 | 2.83 |
| **JUV016** | | | | | | n/a | n/a | 0.52 | 1.35 | 1.72 | 2.22 | 1.31 | 2.21 | 2.24 | 3.19 |
| **JUV017** | | | | | | | | n/a | 0.49 | 0.74 | 1.67 | x | 2.32 | 2.38 | 2.75 |
| **JUV022** | | | | | n/a | 0.67 | x | 1.87 | 1.06 | x | 1.77 | 1.69 | 1.83 | x | |
| **JUV025** | | | n/a | 0.42 | 1.14 | 1.26 | 1.08 | 2.02 | 1.09 | 1.35 | 1.51 | 1.73 | 2.37 | 2.44 | 2.76 |

Sessions included in imaging dataset

- Adult, late
- Adolescent, late
- Adult, early
- Adolescent, early

**Supplemental Table 3. Control analysis using only first 3 sessions after criterion achieved (Figure 3D inset).** Light blue: adult "early" learning sessions. Dark blue: adult "expert" learning sessions. Light pink: adolescent "early" learning sessions. Dark pink: adolescent "expert" learning sessions. Numbers represent overall d' from each session. X indicates imaging or behavioral data unusable for that session.

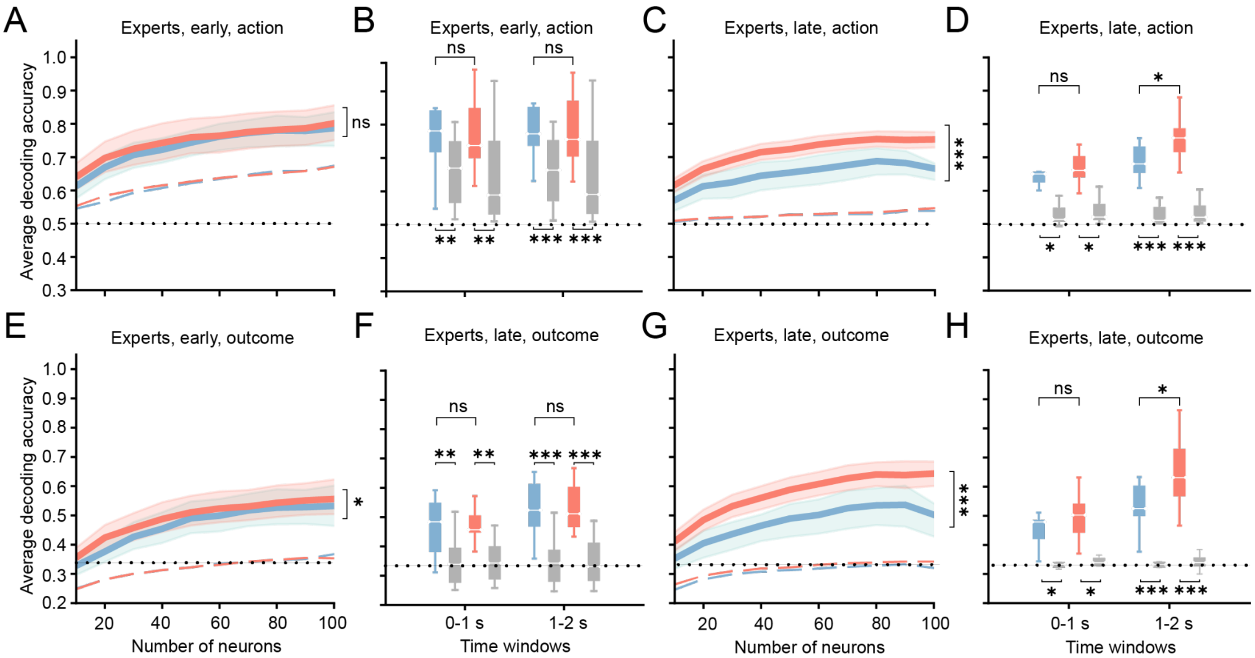


**Supplemental figure 2. Decoding performance of action and outcome in expert adult and adolescent animals. A)** Average decoding accuracy for action between 1-2 s after cue onset as a function of the number of neurons used in the decoding model for expert animals in the early learning stage (two-way ANOVA, no main effect of age, $p$ = 0.646). **B)** Average decoding accuracy of data shown in A) for action binned over 0-1 s and 1-2 s after cue onset. In the early learning phase, data from both age groups performs significantly better than shuffled data, but there was no statistical difference between adult and adolescent decoding performance at either 0-1 s ($p$ = 0.930) or 1-2 s ($p$ = 0.838). Adult data vs. shuffled data, 0-1 s: $p$ = 0.002, 1-2 s: $p$ = 0.002. Adolescent data vs. shuffled data, 0-1 s: $p$ < 0.001, 1-2 s: $p$ < 0.001 (Mann-Whitney U test, FDR_BH corrected)**. C)** Average decoding accuracy for action (as in A) except experts are now in the late learning stage. Two-way ANOVA; main effect of age group, $p$ < 0.001. **D)** In the late learning stage, adolescent difference from adult decoding performance for action neared statistical significance at 0-1 s ($p$ = 0.055), and the difference was significant at 1-2 s ($p$ = 0.014). Adult data vs. shuffled data, 0-1 s: $p$ = 0.016, 1-2 s: $p$ = 0.016. Adolescent data vs. shuffled data, 0-1 s: $p$ < 0.001, 1-2 s: $p$ < 0.001 (Mann-Whitney test, FDR_BH corrected)**. E)** Average decoding accuracy for outcome (Hit, FA, and CR) in the early learning stage of experts. Two-way ANOVA, main effect of age group, $p$ = 0.043. **F)** Same analysis as (B) for experts in the early learning stage for trial type. No statistical difference between adult and adolescent decoding performance at either 0-1 s ($p$ = 1) or 1-2 s ($p$ = 0.977). Adult data vs. shuffled data, 0-1 s: $p$ = 0.002, 1-2 s: $p$ = 0.002. Adolescent data vs. shuffled data, 0-1 s: $p$ < 0.001, 1-2 s: $p$ < 0.001 (Mann-Whitney U test, FDR_BH corrected)**. G)** Average decoding accuracy for outcome (Hit, FA, and CR) in the late learning stage for experts. Two-way ANOVA, main effect of age, $p$ < 0.001. **H)** In the late learning stage, adolescents' difference from adult decoding performance for outcome showed a trend level effect at 0-1 s ($p$ = 0.096) and was significant at 1-2 s ($p$ = 0.042). Adult data vs. shuffled data, 0-1 s: $p$ = 0.016, 1-2 s: $p$ = 0.016. Adolescent data vs. shuffled data, 0-1 s: $p$ < 0.001, 1-2 s: $p$ < 0.001(Mann-Whitney U test, FDR_BH corrected)**.** *p<0.05, **p<0.01, ***p<0.001

**Supplemental Table 4. Average decoding accuracy in 0-1s and 1-2s window after cue onset in a subset of mice that became experts.**

| | | | Data % median±MAD | Shuffle % median±MAD | *p-value* Wilcoxon | *p-value* Mann-Whitney (between ages) |
|---|---|---|---|---|---|---|
| **0-1s** | Early | Adult | 61.6±5.9 | 51.2±1.2 | 0.0273 | 0.977 |
| | | Adol. | 61.6±5.9 | 50.7±1.1 | $2.44 \times 10^{-4}$ | |
| | Late | Adult | 63.7±3.8 | 50.0±0.1 | 0.0156 | 0.0690 |
| | | Adol. | 66.3±4.1 | 50.0±0.2 | $4.77 \times 10^{-4}$ | |
| **1-2s** | Early | Adult | 67.3±4.8 | 50.7±1.0 | 0.0260 | 0.661 |
| | | Adol. | 66.0±5.7 | 50.8±1.4 | $2.44 \times 10^{-4}$ | |
| | Late | Adult | 72.0±4.5 | 50.3±0.5 | 0.0156 | 0.0373 |
| | | Adol. | 82.8±6.6 | 50.0±0.5 | $4.77 \times 10^{-7}$ | |

**Supplemental Table 5. Average decoding accuracy in 1-2s window after cue onset for different trial types in subset of mice that became experts.**

| | | Data % median±MAD | Shuffle % median±MAD | *p-value* Wilcoxon | *p-value* Age, two-way ANOVA |
|---|---|---|---|---|---|
| Hit | Adult | 68.7±7.9 | 48.5±0.4 | 0.0156 | 0.0038 |
| | Adol. | 80.4±5.4 | 48.2±0.6 | $2.38 \times 10^{-7}$ | |
| FA | Adult | 66.3±3.8 | 47.6±0.9 | 0.0156 | 0.0271 |
| | Adol. | 72.8±5.0 | 47.2±0.8 | $2.38 \times 10^{-7}$ | |
| CR | Adult | 75.9±2.9 | 47.4±0.4 | 0.0156 | 0.0051 |
| | Adol. | 79.9±5.6 | 47.5±0.5 | $2.38 \times 10^{-7}$ | |

**Supplemental Table 6. Average decoding accuracy in non-learners.**

| | | Data % **median±MAD** | Shuffle % **median±MAD** | **p-value Wilcoxon** | **p-value Mann-Whitney** |
|---|---|---|---|---|---|
| **0-1s** | **Adult** | 60.0±8.6 | 53.7±2.1 | 0.652 | 0.837 |
| | **Adol.** | 55.9±3.1 | 53.4±3.6 | 0.0313 | |
| **1-2s** | **Adult** | 60.6±6.0 | 53.8±2.3 | 0.568 | 0.758 |
| | **Adol.** | 66.9±4.0 | 53.3±3.7 | 0.0313 | |

**Supplemental Table 7. Average decoding accuracy in 1-2s after cue onset for individual 'go' or 'no-go' pairs.**

| | Cue pairs | Adult % median±MAD | | *p*-value Wilcoxon | Adol. % median±MAD | | *p*-value Wilcoxon |
|---|---|---|---|---|---|---|---|
| | | Data | Shuffle | | Data | Shuffle | |
| **'go'** | 1,2 | 49.3±1.7 | 50.1±0.2 | 1 | 48.9±1.8 | 50.0±0.2 | 0.572 |
| | 1,3 | 49.1±1.7 | 49.5±0.2 | 0.578 | 50.3±1.4 | 50.0±0.3 | 0.547 |
| | 1,4 | 49.0±1.2 | 50.1±0.5 | 0.438 | 49.7±2.5 | 50.1±0.3 | 0.674 |
| | 2,3 | 48.9±1.4 | 49.9±0.2 | 0.938 | 49.2±2.3 | 50.3±0.4 | 0.938 |
| | 2,4 | 48.6±3.0 | 50.0±0.5 | 0.578 | 50.8±1.8 | 50.0±0.8 | 0.388 |
| | 3,4 | 52.7±4.6 | 49.7±0.2 | 0.625 | 48.8±2.4 | 49.9±0.4 | 0.884 |
| **'no-go'** | 5,6 | 50.9±3.7 | 50.1±0.3 | 0.625 | 53.6±2.1 | 50.4±0.8 | $2.55 \times 10^{-3}$ |
| | 5,7 | 50.7±0.9 | 50.1±0.7 | 0.156 | 54.6±2.2 | 50.1±0.5 | $1.05 \times 10^{-4}$ |
| | 5,8 | 51.0±1.5 | 50.5±0.1 | 0.688 | 53.7±3.3 | 50.4±0.8 | $1.63 \times 10^{-3}$ |
| | 6,7 | 49.2±1.3 | 49.9±0.2 | 0.938 | 51.5±1.7 | 50.0±0.4 | $9.66 \times 10^{-3}$ |
| | 6,8 | 49.0±1.3 | 49.8±0.3 | 0.438 | 53.3±1.7 | 50.2±0.5 | $1.01 \times 10^{-3}$ |
| | 7,8 | 51.4±1.4 | 49.5±0.5 | 0.572 | 49.2±2.5 | 49.8±0.3 | 1 |

**Supplemental Table 8. Percentage of variance of the calcium activity explained by running speed (comparing early vs. late)**

| | Adult % median±MAD | Adol. % median±MAD |
|---|---|---|
| **Early** | 19.1±4.1 | 20.3±4.1 |
| **Late** | 25.9±5.2 | 12.8±3.6 |
| ***p*-value Mann-Whitney** | 0.103 | $6.08 \times 10^{-4}$ |

### Supplemental Table 9. Average decoding accuracy in experts with adjusted $\Delta F/F_0$ removing the running effect.

| | | Data % **median±MAD** | Shuffle % **median±MAD** | **p-value Wilcoxon** | **p-value Mann-Whitney** |
|---|---|---|---|---|---|
| **0-1s** | **Adult** | 66.2±3.7 | 50.0±0.2 | 0.016 | 0.158 |
| | **Adol.** | 67.8±3.4 | 50.0±0.3 | $4.77 \times 10^{-7}$ | |
| **1-2s** | **Adult** | 73.5±5.5 | 50.3±0.3 | 0.016 | 0.007 |
| | **Adol.** | 83.5±4.8 | 49.9±0.3 | $4.77 \times 10^{-7}$ | |

### Supplemental Table 10. Average pairwise and population noise correlation

| | | **Adult** mean±SEM | | **Adol.** mean±SEM | |
|---|---|---|---|---|---|
| | | Pairwise | population | Pairwise | Population |
| **Early** | Correct | 0.015±0.002 | 0.308±0.070 | 0.020±0.002 | 0.301±0.073 |
| | Incorrect | 0.014±0.002 | 0.361±0.032 | 0.018±0.002 | 0.341±0.031 |
| **Late** | Correct | 0.023±0.003 | 0.193±0.013 | 0.022±0.002 | 0.309±0.021 |
| | Incorrect | 0.019±0.003 | 0.190±0.017 | 0.022±0.002 | 0.319±0.030 |

### Supplemental Table 11. Average decoding accuracy of real ensemble and pseudo ensemble data.

| | | **Adult %** median±MAD | **Adol. %** median±MAD | *p*-value |
|---|---|---|---|---|
| **Early** | Real | 67.3±4.8 | 66.0±5.7 | 0.661 |
| | Pseudo | 66.6±7.9 | 66.0±4.5 | 0.747 |
| | *p*-value | 0.193 | 0.058 | |
| **Late** | Real | 72.0±4.5 | 82.8±6.6 | 0.037 |
| | Pseudo | 69.6±6.8 | 76.2±5.0 | 0.077 |
| | *p*-value | 0.0156 | $2.10 \times 10^{-5}$ | |
